# Alternative polyadenylation mediates genetic regulation of gene expression

**DOI:** 10.1101/845966

**Authors:** Briana Mittleman, Sebastian Pott, Shane Warland, Tony Zeng, Mayher Kaur, Yoav Gilad, Yang I Li

**Affiliations:** Genetics, Genomics, and Systems Biology, University of Chicago, IL; Department of Human Genetics, University of Chicago, IL; Section of Genetic Medicine, Department of Medicine, University of Chicago, IL

## Abstract

With the exception of mRNA splicing, little is known about co-transcriptional or post-transcriptional regulatory mechanisms that link noncoding variation to variation in organismal traits. To begin addressing this gap, we used 3’ Seq to characterize alternative polyadenylation (APA) in the nuclear and total RNA fractions of 52 HapMap Yoruba lymphoblastoid cell lines, which we have studied extensively in the past. We identified thousands of polyadenylation sites that are differentially detected in nuclear mRNA and whole cell mRNA, and found that APA is an important mediator of genetic effects on gene regulation and complex traits. Specifically, we mapped 602 apaQTLs at 10% FDR, of which 152 were found only in the nuclear fraction. Nuclear-specific apaQTLs are highly enriched in introns and are also often associated with changes in steady-state expression levels, suggesting a widespread mechanism whereby genetic variants decrease mRNA expression levels by increasing usage of intronic PAS. We identified 24 apaQTLs associated with protein expression levels, but not mRNA expression, and found that eQTLs that are not associated with chromatin QTLs are enriched in apaQTLs. These findings support multiple independent pathways through which genetic effects on APA can impact gene regulation. Finally, we found that 19% of apaQTLs were also previously associated with disease. Thus, our work demonstrates that APA links genetic variation to variation in gene expression levels, protein expression levels, and disease risk, and reveals uncharted modes of genetic regulation.

## Introduction

Nearly all genetic variants associated with complex traits are noncoding, suggesting that inter-individual variation in gene regulation plays a dominant role in determining phenotypic outcome. To investigate the function of trait-associated variants identified using genome-wide association studies (GWAS), studies have used regulatory quantitative trait loci (QTL) mapping to associate GWAS loci with variation in mRNA expression levels, DNA methylation levels, and other molecular phenotypes. Although many GWAS loci affect mRNA expression levels (i.e. are eQTLs), several recent discoveries highlight the pressing need for a better understanding of the genetic control of gene regulation, beyond that of just mRNA expression levels. For example, one recent study (Chun et al., 2017) found that the majority of autoimmune GWAS loci do not appear to affect mRNA expression levels. Two other studies observed that many genetic variants affecting variation in protein levels (pQTLs) do not affect mRNA expression levels (Battle et al., 2015; Chick et al., 2016). Altogether these findings indicate that there may be unknown or understudied regulatory mechanisms that link genetic variation to complex traits, and that these mechanisms are independent of changes in the amplitude of mRNA expression levels. Moreover, even when a disease-associated variant is known to impact mRNA expression levels, the mechanisms by which expression is affected is often unclear. Indeed, a third of all eQTLs identified in human lymphoblastoid cell lines (LCLs) are not associated with any chromatin-level regulatory phenotypes including transcription factor binding and histone modifications (Y. I. Li et al., 2016), again raising the possibility that understudied regulatory mechanisms mediate these eQTL effects.

One such understudied mechanism is alternative polyadenylation (APA). Well over half of all human protein coding genes encode multiple polyadenylation sites (PAS), resulting in the production of diverse mRNAs with alternative termination sites (Tian & Manley, 2017; Mayr, 2016; Shi, 2012). Unlike alternative mRNA splicing, which leads to changes in splice site selection, APA leads to changes in the transcript termination site, often resulting in 3’ untranslated regions (UTRs) with different lengths. As 3’UTRs are densely packed with regulatory elements that impact mRNA stability, miRNA binding, and mRNA localization (reviewed in (Mayr, 2017; Tian & Manley, 2017)), genetic control of APA may be a key mechanism by which genetic variants impact gene regulation, including mRNA expression levels, without affecting chromatin-level phenotypes such as promoter or enhancer activity. Moreover, proteins translated from different APA isoforms may differ in length and protein-protein interactions, and these differences can have phenotypic effects. For example, global increased usage of intronic PAS has been shown to increase risk for multiple myeloma and chronic lymphocytic leukemia (Lee et al., 2018; Singh et al., 2018) by producing mRNAs that translate into truncated proteins, which causes defects in tumour-suppressive functions (Lee et al., 2018; Singh et al., 2018).

To evaluate the role of APA in mediating genetic effects on gene expression and disease, we sought to identify genetic variants associated with APA on a genome-wide scale. To date, the few studies that have used genome-wide methods to identify variants associated with APA (apaQTLs) have used existing RNA-seq data to infer PAS locations and usage (L. Li, Gao, Peng, Wagner, & Li, 2019; Yoon, Hsu, Im, & Brem, 2012; Yang et al., 2019; Bonder et al., 2019). While using existing RNA-seq to study APA is economical, identifying PAS and estimating usage using RNA-seq are error-prone and often imprecise (Ha, Blencowe, & Morris, 2018). Furthermore, using existing whole-cell, total RNA-seq data is not informative with regards to whether inter-individual differences in PAS usages are the result of variation in transcriptional termination site choice, or isoform-specific decay or export. Here, we used 3’ RNA-seq (3’ Seq) to measure PAS usage in steady-state mRNA collected from whole cells as well as mRNA collected from the nucleus, which is comprised of a high proportion of nascent mRNA. This design allowed us to study the effect of genetic variation on isoform PAS at multiple stages of the mRNA lifecycle. Importantly, we collected these data from a panel of human lymphoblastoid cell lines (LCLs) that were previously profiled in great molecular detail, including measurements at the chromatin, RNA, and protein levels (Degner et al., 2012; McVicker et al., 2013; Y. I. Li et al., 2016; Pickrell et al., 2010). Integrating the apaQTLs we identified with previously collected data types allowed us to characterize the functional impact of variation in APA on each of the major steps of the gene regulatory cascade. We use these data to show that genetic effects on polyadenylation can independently affect virtually all steps of gene regulation (mRNA expression level, translation rate, and protein expression level), and that such effects can be associated with protein expression, but not RNA expression.

## Results

To measure the impact of inter-individual variation in APA on multiple stages of gene regulation, we quantified PAS usage in a panel of 52 Yoruba HapMap LCLs. These same samples have been the subjects of multiple studies of gene regulation over the last decade (Degner et al., 2012; McVicker et al., 2013; Y. I. Li et al., 2016; Pickrell et al., 2010). We applied 3’ Seq to mRNA collected from whole cells (total fraction) of 52 LCLs to comprehensively identify PAS and estimate usage without relying on existing annotations. In addition, to capture polyadenylated mRNA that may be under-represented or absent in the total fraction due to rapid turnover, we separately applied 3’ Seq to mRNA from isolated nuclei (nuclear fraction) of the same 52 LCLs (Fig. 1A).

**Figure 1:**
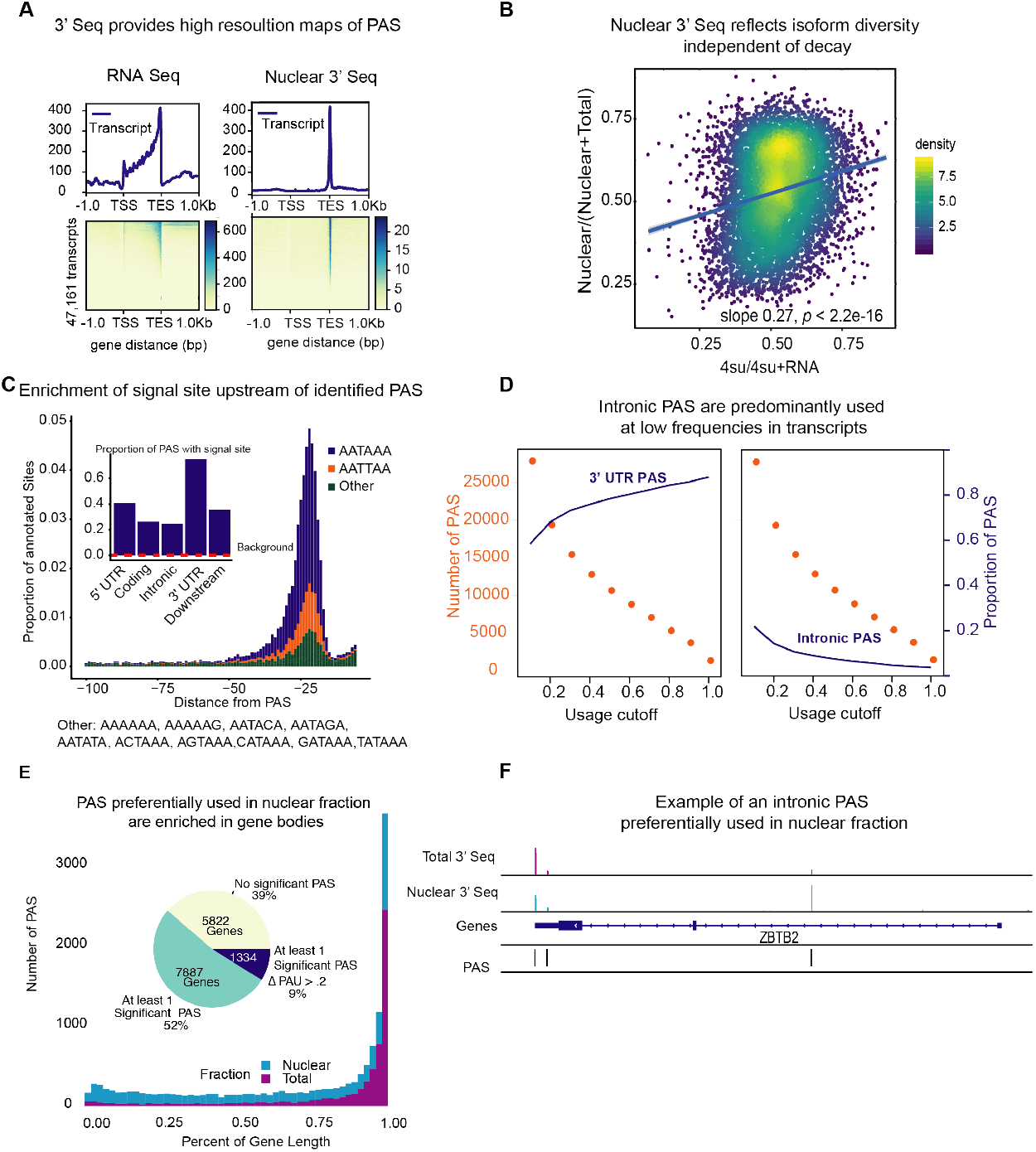
**(A)** Meta gene plot showing read coverage for five RNA sequencing libraries collected from LCLs (Pick-rell) *(Left)* and for five 3’ Seq libraries collected from nuclei isolated from LCLs *(Right)*. **(B)** Nuclear 3’ Seq capture polyadenylation of nascent transcripts. The ratio of new mRNA to steady-state mRNA (Y. I. Li et al., 2016) (x axis) are plotted against the ratio of 3’ Seq reads from the nuclear fraction to 3’ Seq reads from the total fraction (y axis). **(C)** *(Main)* Density of canonical (AATAAA, AATTAA) and other polyadenylation signal sites upstream of identified PAS. *(Inset)* Proportion of PAS in different genomic regions with a polyadenylation signal site 10–50bp upstream of cleavage site. The red dotted line represents the proportion of signal site in random 40bp windows, i.e. the intronic background. **(D)** Number (orange) and proportion (purple) of PAS in the 3’ UTR (*Left*) and introns (*Right*) are plotted against usage cutoff in the nuclear fraction. The proportion of intronic PAS increases as the usage cutoff decreases, implying that a disproportionate number of intronic PAS are used at low frequencies. **(E)** *(Main)* Meta gene plot showing the number of differentially used PAS identified by LeafCutter (Online Methods) with a Δ PAU of 0.20 across the gene body. *(Inset)* Estimated number of genes identified with differential PAS usage between total and nuclear fractions. **(F)** *ZBTB2* was identified to harbor a differentially used PAS between total and nuclear fractions. 3’ Seq tracks represent aggregated read counts from all 52 individuals.

### Nuclear 3’ Seq measures PAS usage independent of post-transcriptional decay

We first verified that the nuclear 3’ Seq data capture transcripts at a more primitive stage compared to the total 3’ Seq data, and thus better reflect mRNA diversity independent of decay. To this end, we reasoned that genes with higher nuclear 3’ Seq read counts relative to total 3’ Seq read counts should show faster decay on average. Indeed, we found that the relative number of nuclear 3’ Seq reads to total 3’ Seq reads is positively correlated with both the ratio of 4sU-seq to RNA-seq read counts (Y. I. Li et al., 2016) (*p* < 2.2^−16^, Fig. 1B) and direct gene level mRNA decay estimates (Pai et al., 2012) (*p* < 2.2^−16^, Supplementary Fig. 1), two different measures of decay. After filtering the 3’ Seq data for possible internal priming (Methods), we identified 41,810 PAS in 15,043 genes. We found that 67% of the protein coding genes expressed in LCLs harbor multiple PAS, suggesting that APA can impact the regulation of most genes (Tian & Manley, 2017; Mayr, 2016; Shi, 2012). We found that the polyA binding protein motif (AATAAA), also known as the polyadenylation signal site, is the most strongly enriched protein binding motif in regions surrounding our PAS (*p* < 10^−391^). We observed that PAS in the 3’ UTR are more likely to have a polyadenylation signal compared with intronic PAS (*p* < 10^−16^, difference of proportion t-test, 75.0% vs 24.8%,) (Fig. 1C, Supplementary Fig. 3) and that nearly half (48.3%) of all 41,810 PAS we identified are located in 3’ UTRs (19.4× enrichment) (Singh et al., 2018). Nevertheless, despite an overall depletion of PAS in introns (0.35× genome-wide levels), we found that the number of PAS in introns is notable (12,793/41,810; 30.6%) (Fig. 1D, Supplementary Fig. 2). While signal sites were more highly enriched near 3’ UTR PAS than intronic PAS, PAS in introns show clear enrichment of polyadenylation motif 10-50 bp upstream of the cleavage site compared to background intronic sequences (24.8% vs 0.24% *p* < 10^−16^, difference of proportion t-test, Fig. 1D). Thus, the recognition of intronic polyadenylation signals is a general mechanism that can result in premature termination of transcription. In addition, although slightly enriched in the first introns of genes (2.69× enrichment over uniform distribution), intronic PAS could be identified in all introns and thus are not likely to result from defective telescripting activity alone (e.g. from a depletion in U1 small nuclear ribonucleoprotein (snRNP)) (Kaida et al., 2010; Berg et al., 2012; Oh et al., 2017).

We also observed that intronic PAS have on average lower usage across individuals than PAS located in 3’ UTRs (16.9% vs 46.2%). These differences may be explained by weaker polyadenylation signals at intronic PAS compared to 3’ UTR PAS, but we hypothesized that some intronic PAS might have low usage because premature polyadenylation at intronic sites can produce short-lived transcripts that are rapidly degraded and thus under-represented in the total mRNA fraction. To test this hypothesis, we identified PAS that are used more often, or exclusively, in the nuclear fraction compared to the total fraction. By comparing PAS usage estimated in the nuclear and total fractions from all 52 individuals, we identified 591 PAS in 585 genes that are used more often in the nuclear fraction (10% FDR). 134 of these 591 PAS were found to be used by 1% or less of the transcripts in the total fraction, suggesting that these transcripts may be absent from the cytoplasm (Fig. 1E, Supplementary Fig. 4, Methods). Notably, we found that 387 of the nuclear-enriched PAS are intronic (Supplementary Fig. 4; see example in Fig. 1F), a large proportion of which (83.4% vs 43% for all PAS) are absent from a comprehensive annotation of PAS compiled from 78 human studies that used 3’ Seq (Methods, Supplementary Fig. 5) (Wang, Nambiar, Zheng, & Tian, 2018). These findings suggest that mRNA transcripts are polyadenylated in introns at a higher frequency than generally appreciated, and that many of these isoforms escape detection from studies of total mRNA owing to their rapid decay.

### Genetic loci associated with variation in APA

We next sought to identify genetic loci associated with inter-individual variation in APA. We quantified APA as the normalized ratio of reads mapping to each PAS compared to reads mapping to all PAS assigned to the same gene (Methods, Supplementary Fig. 6-7). We tested cis-association between genetic variants and PAS usage, correcting for batch and the top principal components (Methods, Supplementary Fig. 8). Using 3’ Seq data from the nuclear fraction, we identified 602 nuclear apaQTLs in 479 genes at 10% FDR. In the total fraction, we identified 443 apaQTLs in 353 genes at 10% FDR. For example, individuals with the C/C genotype (rs11032578) are more likely to use an intronic PAS in the *ABTB2* gene compared to individuals that are heterozygous C/T or homozygous T/T (Fig. 2A). In both fractions, apaQTLs occur near the PAS they most strongly correlate with and are located at the 3’ ends of gene bodies (Fig. 2B-C). While the proximity of the apaQTLs to PAS may suggest that genetic variants that affect polyadenylation signal motifs drive most of the genetic effects on APA, we found limited evidence that supports this possibility (Supplementary Fig. 9).

**Figure 2:**
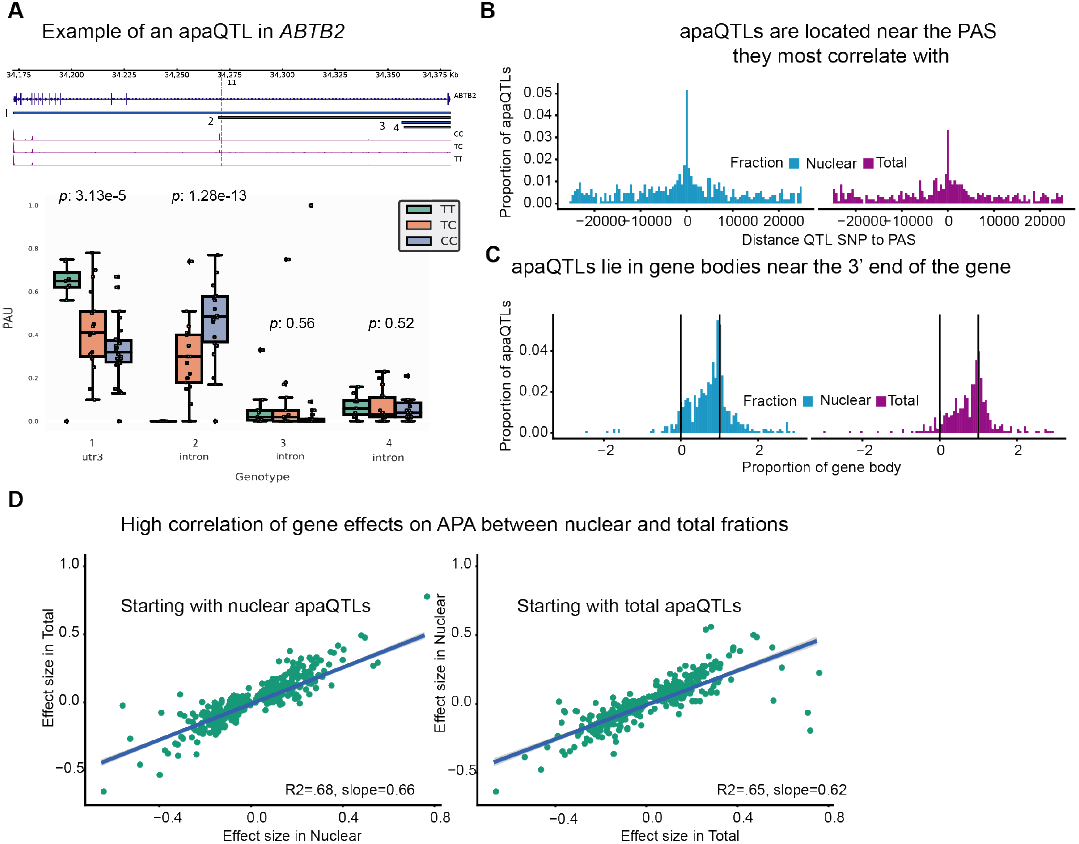
**(A)** An apaQTL in the *ABTB2* gene impact usage of an intronic PAS. *(Top)* Gene track and identified PAS. Each bar represents a potential isoform. The red bar corresponds to the isoform most strongly associated with the apaQTL. The vertical dotted line represents the position of the strongest apaQTL SNP. *(Bottom)* Polyadenylation site usage at each PAS by genotype listed according to isoform order above. The C allele increases usage of the intronic PAS. **(B)** Location of the top nuclear (*Left*) and total (*Right*) apaQTL SNPs relative to their corresponding PAS. **(C)** Meta gene plot showing the distribution of apaQTL SNPs in the annotated gene body, where 0 represents the TSS and 1 represents the annotated transcription end site. **(D)** Effect sizes of apaQTLs originally identified at 10% FDR in nuclear *(Left)* and total *(Right)* fraction plotted against the effect sizes ascertained in total and nuclear fractions, respectively.

Next, we quantified the sharing and specificity of genetic effects on APA in the nuclear and total fractions. We estimated that the vast majority of nuclear apaQTLs were shared with total apaQTLs (*π*_1_ 1=0.85) and vice-versa (*π*_1_=0.87, Supplementary Fig. 10). Additionally, we observed that their effect sizes were highly correlated (*r*^2^ = 0.66; *p* = 10^−16^, Fig. 2D, Supplementary Fig. 11). These results suggest that the predominant mechanism by which genetic variants affect steady-state PAS usage is to directly impact PAS choice rather than to affect the stability of an isoform ending at one site relative to that with another ending (e.g. by affecting isoform-specific decay). Nevertheless, we identified 153 nuclear-specific apaQTLs and 97 total-specific apaQTLs (Methods).

### APA explain eQTLs that are not associated with chromatin phenotypes

Given that most apaQTLs identified in our study are represented in the nuclear fraction, we focused on the 602 nuclear apaQTLs for subsequent analyses. Although APA produces isoforms with distinct ends, it is possible that the isoforms are functionally identical, especially when they differ only in 3’ UTR lengths and thus encode the same protein sequence. To better understand the functional impact of apaQTLs, we asked whether they are also associated with changes in gene or protein expression levels. We found that genes associated with apaQTLs are enriched for both genes with eQTLs (eGenes, Wilcoxon rank sum test *p* = 1.36 × 10^−12^) and genes with protein expression QTLs (pGenes; *p* = 0.0006) compared to genes with neither association (Fig. 3A, Supplementary Fig. 12) (Y. I. Li et al., 2016; Battle et al., 2015). Notably, we found that nuclear-specific apaQTLs are even more enriched for eGenes (*p* = 0.002) compared to apaQTLs that are shared in both fractions. This observation led us to hypothesize that intronic apaQTLs affect gene expression levels by increasing the number of transcripts that use premature intronic PAS, of which many may be subject to rapid decay. Indeed, we found a negative correlation between the genetic effect sizes for intronic PAS usage and mRNA expression levels (*p* = 8.97 × 10^−7^, Fig. 3B, Supplementary Fig. 13). Thus, our analyses suggest a widespread mechanism whereby genetic variants decrease mRNA expression levels by increasing usage of premature PAS located in introns. Of note, 13 of the apaQTLs that were detected only in the nuclear fraction are also eQTLs, which highlights the importance of considering multiple stages of mRNA biogenesis to uncover eQTL mechanisms.

**Figure 3:**
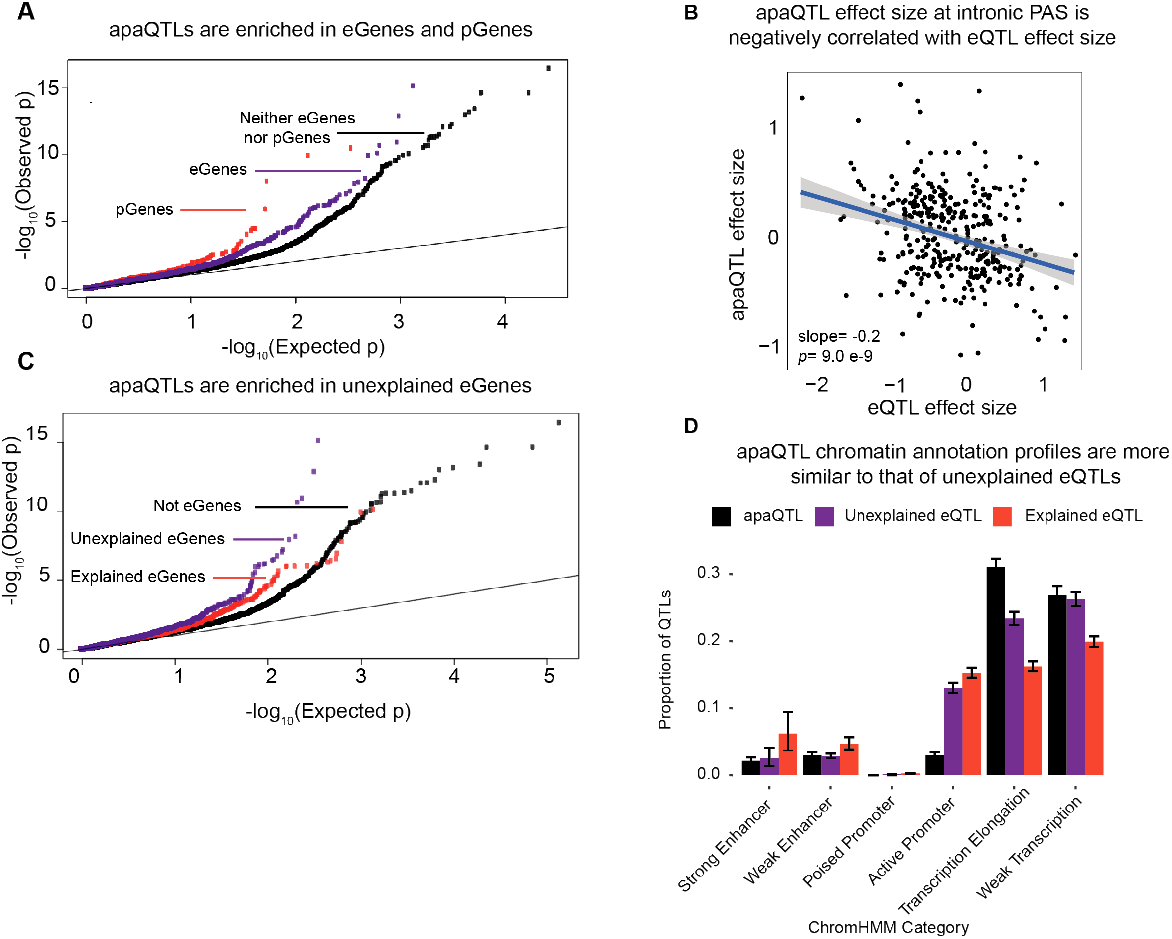
**(A)** Quantile-quantile (Q-Q) plot for apaQTLs shows an enrichment in both eGenes and pGenes. **(B)** Scatter plot of intronic apaQTL effect sizes plotted against their eQTL effect sizes shows negative correlation. **(C)** Quantile-quantile (Q-Q) plot for apaQTLs shows that apaQTLs are more highly enriched in unexplained eGenes compared to explained eGenes. **(D)** Proportion of apaQTL, explained eQTL, and unexplained eQTL SNPs in different genomic annotations. The annotation profiles of apaQTLs show more similarity to that of unexplained eQTLs than to that of explained eQTLs. Error bars represent 95% confidence intervals from bootstraps (Online Methods).

To further investigate the contribution of APA to gene expression, we focused on a set of eQTLs that we previously classified as those with explained putative mechanisms eQTLs (1164 eQTLs, ~ 60%) or as unexplained eQTLs (801 eQTLs, ~ 40%) using data from the same LCLs (Y. I. Li et al., 2016). The eQTLs with explained putative mechanisms were associated with chromatin-level phenotypes including DNase-I hypersensitivity, histone marks, or DNA methylation, and thus are likely to be mechanistically explained by effects mediated by chromatin-level phenotypes (e.g. enhancer or promoter activity). By contrast, 40% of eQTLs were not associated with any chromatin-level measures, and thus their mechanisms of action remain unknown. To test whether apaQTLs might account for unexplained eQTLs, we first asked whether genes with unexplained eQTLs were more likely to also harbor apaQTLs than compared to genes with explained eQTLs. Indeed, we found a significantly higher enrichment of low p-value associations with APA for genes with unexplained eQTLs (*p* = 0.01) (Fig. 3C, Supplementary Fig. 14). We also found that apaQTLs exhibited a chromatin enrichment profile that was more similar to the unexplained eQTLs than the explained eQTLs. In particular, apaQTLs and unexplained eQTLs were more likely to lie in regions of transcription elongation or are associated with weak transcription, and less likely to lie in enhancers or promoters than explained eQTLs (Fig. 3D. Overall, we estimated that the apaQTLs can provide a putative mechanism for 17.3% of otherwise unexplained eQTLs (see Methods). For example, an unexplained eQTL for *C10orf88* (rs7904973) colocalizes with an apaQTL associated with increased usage of an intronic PAS (Fig. 4A). This observation thus highlights APA as one important mechanism by which genetic variation impacts gene expression without affecting enhancer and promoter activity.

**Figure 4:**
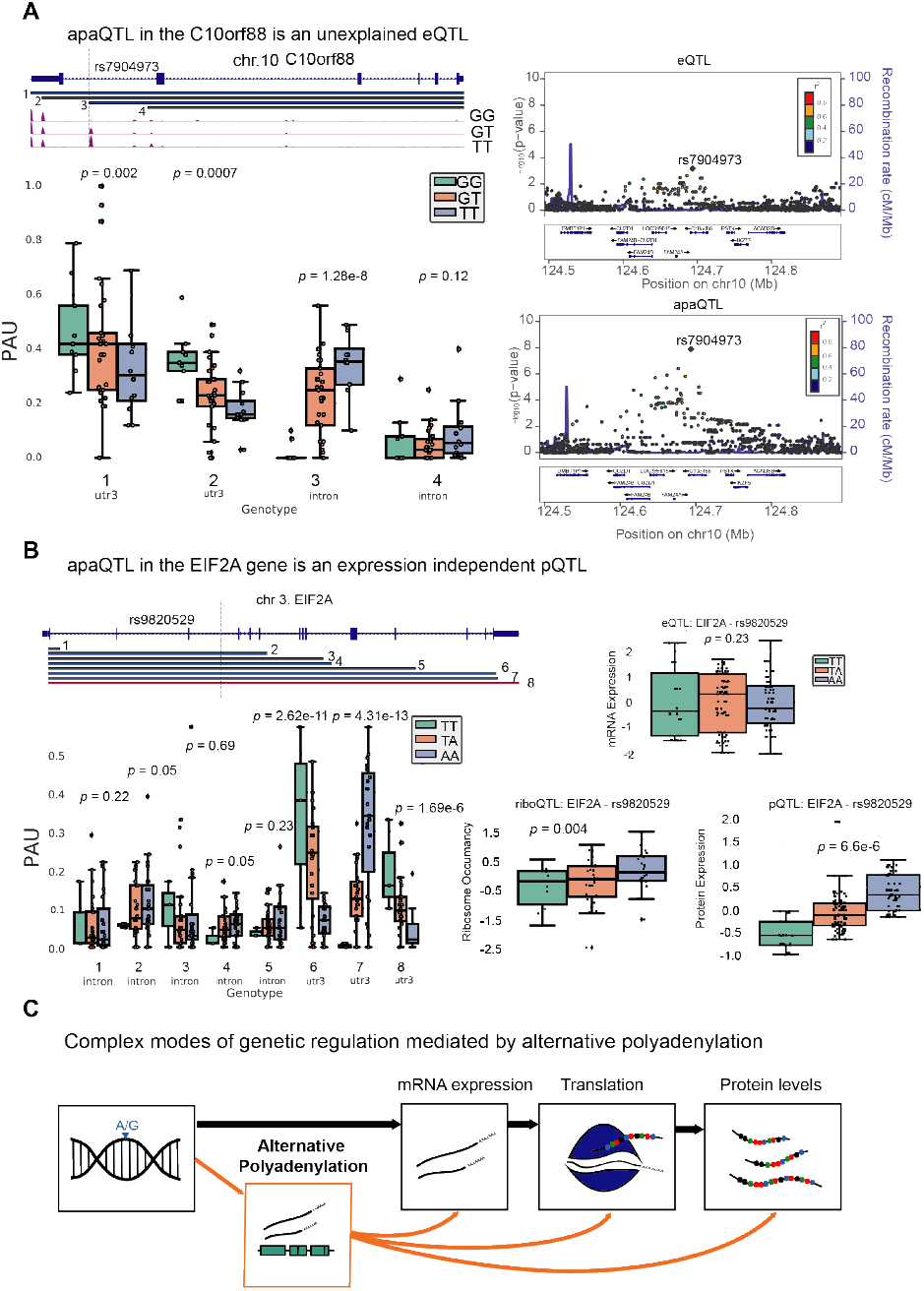
**(A)** Example of an apaQTL that is also an unexplained eQTL in *C10orf88*. *(Top Left)* Gene track and identified PAS in the *C10orf88* gene. The red bar corresponds to the isoform most strongly associated with the apaQTL. The vertical dotted line represents the position of the strongest apaQTL SNP. *(Bottom Left)* Polyadenylation site usage (PAU) at each PAS by genotype listed according to isoform order above. (*Right)* Locus zoom plot for eQTL and apaQTL associations. Interestingly, the top associated SNP, rs7904973, has been linked to increased LDL cholesterol through GWAS (Klarin et al., 2018). **(B)** *(Top Left)* Gene track and identified PAS in the *EIF2A* gene. *(Bottom Left)* Polyadenylation site usage (PAU) at each PAS by genotype listed according to isoform order above. *(Right)* Boxplots showing normalized mRNA expression, ribosome occupancy, and protein expression for *EIF2A* by genotype at the apaQTL SNP (rs9820529). **(C)** Diagram representing multiple pathways by which genetic variation mediates gene regulation through alternative polyadenylation.

### APA mediates gene regulation independently of mRNA expression levels

Previous joint analyses of molecular QTLs suggested that functional genetic variants tend to affect gene regulation in a simple and straightforward manner: first impacting chromatin activity, then mRNA expression, and finally protein expression (Y. I. Li et al., 2016; Battle et al., 2015). However, we found 24 apaQTLs that affect protein expression, but not mRNA expression (Supplementary Table 1), suggesting a more complex mode of gene regulation independent of mRNA expression that involves APA. We found that five of these 24 apaQTLs were significantly associated with ribosome occupancy (Supplementary Table 1). This finding is particularly noteworthy because nearly all genetic effects on ribosome occupancy have been proposed to be mediated by effects on mRNA expression (Battle et al., 2015). Yet, here we provide direct evidence that APA can mediate genetic effects on ribosome occupancy without affecting mRNA expression levels. For example, the apaQTL in the *EIF2A* gene that is associated with a switched usage of two 3’ UTR PAS, colocalizes with a pQTL and a ribosome occupancy QTL (Fig 4B, Supplementary Fig. 15), but is not associated with *EIF2A* mRNA levels (Fig. 4B). Interestingly, the QTL in *EIF2A* affects usage of two PAS in the same 3’ UTR implying that the protein sequence encoded by the two isoforms are identical. Thus, the regulatory associations uncovered at *EIF2A* cannot simply be explained by differences in protein isoform stability. Moreover, while differences in 3’ UTR are often assumed to play a regulatory function by influencing decay (Mayr, 2017), mechanisms involving RNA decay cannot be operational in this case because steady-state mRNA expression is unchanged. Instead, differences between the two isoforms may reflect differential binding of factors that impact translation (Yamashita & Takeuchi, 2017), or differential rates of translation re-initiation at the end of a translation cycle (Rogers, Böttcher, Traulsen, & Greig, 2017).

We identified 19 pQTLs that were associated with APA but not steady-state gene expression or ribosome occupancy levels. Two previous studies also reported the discovery of pQTLs that were not eQTLs. In both studies, the authors proposed that some genetic effects on protein expression levels were mediated by changes in the protein sequence, which would manifest post-translationally. Our finding reveals yet another complex mode of genetic regulation of protein expression level by APA (e.g. perhaps by affecting recruitment of interacting proteins). Thus, these findings provide clear evidence that APA can affect protein expression levels without affecting gene expression levels through multiple regulatory pathways.

### APA mediates genetic effects on complex traits

Lastly, we hypothesized that genetic variation may impact disease risk through APA. Indeed, 19.3% of apaQTLs (including SNPs in LD) are significantly associated with at least one trait in the UCSC GWAS catalog (Kent et al., 2002). Interestingly, an apaQTL in the C10orf88 gene (rs7904973) has been associated with increased LDL cholesterol (Klarin et al., 2018), suggesting that eQTLs mediated by APA can impact organismal phenotypes. APA is complex regulatory mechanism relevant to our understanding of how genetic variation can affect disease; therefore, comprehensive maps of apaQTLs can enhance our ability to interpret GWAS loci, particularly when the implicated variants are not eQTLs (Joehanes et al., 2017; Lee et al., 2018). For example, an apaQTL in the *ELL2* gene (rs56219066) is correlated with increased usage of an intronic PAS and is associated with risk for multiple myeloma. (Swaminathan et al., 2015) Interestingly, multiple myeloma is among the cancer types in which widespread dysregulation of intronic APA has been documented previously (Singh et al., 2018; Lee et al., 2018).

## Discussion

Obtaining a comprehensive understanding of the mechanisms that affect gene regulation is crucial for the functional interpretation of noncoding genetic variation. Yet, existing studies that examine the role of genetic variation on APA are generally characterized by two important shortcomings. Firstly, the study of inter-individual variation in PAS usage have been mostly restricted to APA in the 3’ UTRs (L. Li et al., 2019; Yoon et al., 2012; Yang et al., 2019), leaving genetic variants that impact PAS usage in other regions, e.g. intronic PAS, understudied. Secondly, nearly all existing studies use standard RNA-seq to estimate PAS usage, which not only limits the accuracy of usage quantification, but also makes it difficult to disentangle the contribution of co-transcriptional mechanisms to APA regulation from post-transcriptional mechanisms such as isoform-specific decay.

To overcome these shortcomings, we applied 3’ Seq to total and nuclear cell fractions separately to directly measure PAS usage including that of PAS in intronic regions. These data allowed us to study the effects of genetic variation on polyadenylation at multiple stages of the mRNA life cycle, and to distinguish putative regulatory mechanisms by noting the stages at which the genetic effects on APA were observed. For example, genetic variants can impact steady-state isoform ratio either co-transcriptionally by affecting PAS choice during transcription, or post-transcriptionally by affecting binding of miRNAs or RNA-binding proteins and consequently isoform decay. However, we found that the vast majority of genetic variants that affect PAS usage ratio in total mRNA, were also found to have highly similar effect sizes on PAS usage ratio in the nucleus. This observation implies that inter-individual variation in steady-state APA levels can generally be explained by variation in co-transcriptional mRNA processing, or mRNA processing that occur very soon after transcription.

There are several co-transcriptional mechanisms that may result in variation in PAS usage. For example, previous reports have suggested that variation in the polyadenylation signal site may cause variation in PAS usage. While we found that this was the case for a handful of examples, disruption of canonical signal motifs does not appear to be a major mechanism for generating apaQTLs, an observation that is also supported by a recent study on APA in GTEx data (Supplementary Fig. 9) (L. Li et al., 2019). Other possible co-transcriptional mechanisms involved in PAS choice include competition between the spliceosome and polyadenylation factors for example mediated by the spliceosomal RNA U1 (Oh et al., 2017), and RNAP II pausing (Fusby et al., 2016). Indeed, recent studies have reported that sequence and chromatin context can pause or slow down RNAP II elongation across the gene body (Mayer et al., 2015), suggesting that variation in RNAP II pausing may impact PAS choice (Fusby et al., 2016). For example, in Drosophila melanogaster, paused RNAP II promotes the recruitment of ELAV on the pre-mRNA, which prevents usage of a proximal PAS (Oktaba et al., 2015). Interestingly, the human ortholog, ELAVL1 has been implicated in mRNA localization and may influence APA through competition for binding with other factors (Neve, Patel, Wang, Louey, & Furger, 2017; Berkovits & Mayr, 2015; Dai, Zhang, & Makeyev, 2012).

Although our data suggest that apaQTLs do not generally impact rates of mRNA decay, e.g. by affecting miRNA or RBP binding motifs, we found clear evidence that apaQTLs may promote polyadenylation site choices that result in the production of isoforms with different rates of decay. For example, we observed that genetic variants that increase the usage of isoforms ending at intronic PAS tend to be associated with lower levels of gene expression. This observation is consistent with reports that isoforms with premature polyadenylation are often substrates for nonsense mediated decay or nonstop decay (Tian & Manley, 2017; Vasudevan, Peltz, & Wilusz, 2002). More generally, our results suggest that apaQTLs can affect gene expression levels post-transcriptionally by impacting the production of isoforms with varying levels of stability. Overall, our study highlights APA as an eQTL mechanism independent of promoters and enhancers.

While the effect of genetic variants on gene regulation is generally assumed to move linearly from chromatin, to mRNA, to protein level, our findings reveal several complex modes of genetic regulation for both gene expression and protein expression levels by APA (Fig. 4C). Although we were unable to study the genome-wide effects of APA on protein expression owing to a scarcity of protein-level data, we identified several apaQTLs that affect protein, but not gene expression levels. These results strongly suggest that APA can affect protein expression levels without affecting gene expression levels, because our power to detect genetic effects on gene expression levels far exceeds that to detect genetic effects on protein expression levels. Furthermore, some of these pQTLs were associated with ribosomal occupancy and some were not, which implies multiple pathways by which genetic variants can impact protein expression levels through APA.

In conclusion, there are many pathways through which genetic variants can impact gene regulation and, consequently, organismal phenotypes. While many studies have demonstrated the importance of gene expression regulation through promoters or enhancers, very few studies have focused on co- or post-transcriptional gene regulation. Our study shows that co- and post-transcriptional processes such as APA can mediate the effects of a substantial number of genetic variants on mRNA expression levels, protein expression levels, and risk for complex diseases.

## Methods

### Cell Culture

We cultured 54 Epstein-Barr virus transformed LCLs under identical conditions at 37 C and 5% CO2. These LCLs were derived from Yoruba individuals originally collected as part of the HapMap project (International HapMap Consortium, 2005; Moll, Ante, Seitz, & Reda, 2014). Details for each cell line are found in Supplementary Table 2. We grew cells in a glutamine depleted RPMI [RPMI 1640 1× from Corning (15-040-CM)] completed with 15% FBS, 2mM GlutaMAX (from gibco (35050-061), 100 IU/ml Penicillin, and 100 ug/ml Streptomycin. After passaging them 3 times the lines were maintained at a concentration of 1 × 10^6^ cells per mL. In preparation for extraction, we allowed the cells to grow until a concentration of 1 × 10^6^ cells per mL was reached and then proceeded to extraction.

### Collection and RNA extraction

We collected 30 million cells from each line and divided them into two 15 million cell aliquots. We spun the cells down at 500 RPM at 4C for 2 min, and then washed the pellets with phosphate-buffered saline (PBS) and spun down again. After this we aspirated the PBS, leaving the cell pellet. All washing steps occurred on ice or in cooled centrifuges. At this point every cell line had two separate pellets each from an input of 15 million cells. From each line we took one of these pellets for nuclear isolation. We then carried out nuclear isolation using the nuclear isolation steps outlined by (Mayer & Churchman, 2016). Once we washed and spun down the pellets in the nuclei wash buffer, we resuspended them in 700 ul of the QIAzol lysis reagent (Qiagen). We extracted both RNA cell pellets from the same line in the same batch using the miRNeasy kit (Qiagen) according to manufacture instructions, including the DNase step to remove potentially contaminated genomic DNA. Details for the collection such as cell viability and cell concentration at time of collection are found in Supplementary Table 2. We checked the quality of the collected RNA using a nanodrop. RNA concentrations and absorbance levels from the collection are in Supplementary Table 2.

In order to verify fraction separation, we completed the Mayer and Churchman protocol to isolate chromatin and collected cell lysates for each step in the fractionation (Mayer & Churchman, 2016). We performed western blots against both GAPDH (GAPDH antibody (6C5) Life Technologies AM4300) and the Carboxyl Terminal Domain of Pol-II (CTD) (Pol II CTD Ser5-P antibody, Active Motif, 61085). We ran each lysate on Mini-protean TGX precast gels (bioRad 456-1093) after digesting any remaining DNA molecules from the nuclear isolate with benzonase nuclease. We used Goat anti-Mouse IgG (H+L) (Invitrogen 32430) as a secondary antibody for the GAPDH antibody and Goat anti-Rat IgG (H+L) (Invitrogen 31470) as a secondary antibody for the CTD antibody. We diluted all antibodies in a 1:1000 dilution with blocking solution made from dry milk (LabScientific Lot 1267N Cat M0841). We show GAPDH isolated in the cytoplasm and CTD to the chromatin fraction (Supplementary Fig. 16).

### 3’ Sequencing library generation

We generated 108 single-end RNA 3’ sequencing libraries from the total and nuclear RNA extract using the QuantSeq 3’ mRNA-Seq Library Prep Kit (Moll et al., 2014) as directed by the manufacturer. We used 5ng of each sample as input. We submitted the libraries for sequencing on the Illumina NextSeq5000 at the University of Chicago Genomics Core facility using single end 50bp sequencing.

### 3’ Sequencing data processing

We mapped 3’ Seq reads to hg19 (Church et al., 2011) using STAR RNA-seq aligner (Dobin et al., 2013) using default settings with the WASP mode to filter out reads mapping with allelic bias (van de Geijn, McVicker, Gilad, & Pritchard, 2015). Similar to previously published 3’ Seq methods, we accounted for internal priming by filtering reads preceded by 6 Ts in a row or 7 of 10 Ts in the 10 bases directly upstream of the mapping position in the reference genome (Tian, Hu, Zhang, & Lutz, 2005; Sheppard, Lawson, & Zhu, 2013; Beaudoing, Freier, Wyatt, Claverie, & Gautheret, 2000). We verified the individual identity of all bam files using VerifyBamID (Jun et al., 2012). Due to low confidence in the identity of 2 individuals, they were removed from all analysis. Raw read and mapped read statistics after accounting for internal priming can be found in Supplementary Table 1 (Supplementary Fig. 17).

### Identification of PAS

We merged all mapped reads and called peaks using an inclusive method, identifying all regions of the genome with non-zero read counts in 90% percent of libraries and an average read count of greater than 2 counts. This resulted in 138,181 peaks. We assigned each of these peaks to a genic location according to NCBI Refseq annotations for 5’ UTRS, 3’ UTRs, exons, introns, and regions 5kb downstream of annotated genes downloaded from the UCSC table browser (Kent et al., 2002). When a region mapped to multiple genes we used a hierarchical model, similar to the method used by Lin et al. (Lin et al., 2012) to assign the peak to a gene annotations. Our method prioritizes annotations in the following order: 3’ UTRs, 5kb downstream of genes, exons, 5’ UTRs, and introns. To further verify absence of PAS detected as a result of internal priming we removed PAS with 6A’s or 70% As in the 15 basepairs downstream of the site. We next utilized a gene level noise filter to account for non-uniform read coverage across the genome. We created a usage score for each PAS based on of the number of reads mapping to the PAS over the number of reads mapping to any PAS associated with the same gene. We filtered out peaks with a mean usage of less than 5% in both the total and nuclear libraries. After this filter, we were left with 35,032 PAS in the total fraction and 39,164 PAS in the nuclear fraction. The merged set with PAS from both fractions used for PAS QC is available on GEO and has 41,810 PAS. We compared our set of PAS to the human PolyADB release 3.2 annotation (Wang et al., 2018)(Supplementary Fig. 5).

### PAS Signal site enrichment and locations

To explore the location of the signal site relative to the PAS (most 3’ end of each identified peak), we determined the relative position of previously described potential signal sites to this position (Beaudoing et al., 2000). We then extended each PAS 100bp upstream and identified the starting position of each of the 12 PAS signal site variations identified by Beaudoing et al. without allowing for sequence mismatch (Beaudoing et al., 2000).

### Differential Isoform analysis

We mapped 3’ Seq reads to all PAS peaks with mean coverage of 5% in the total or nuclear fraction libraries. This results in 41,813 annotated sites. We assigned reads to PAS using the featureCounts tool with the -O flag to assign reads to all overlapping features (Liao, Smyth, & Shi, 2014). We ran the leafcutter ds.R script on chromosomes 1-22 separately using the cellular fraction label as the sample group identifier (Y. I. Li et al., 2018). This analysis tests 9790 genes and resulted in 8227 genes with significant (FDR 10%) isoform level differences between the total and nuclear cellular fraction. We called differentially used PAS as sites with a Δ polyadenylation site usage (Δ PAU) greater than 0.2 or less than −0.2. In our analysis a positive ΔPAU corresponds to increase usage in the total cellular fraction while a negative Δ PAU corresponds to increased usage in the nuclear fraction.

### Relationship with nascent transcription

We used 30 min 4su data and RNA decay measurements collected in the same panel of LCLs as used in this study. RNA decay data was originally collected and processed in Pai et al. 2012. (Pai et al., 2012) The 4su data collection and processing can be found in Li et al. 2016(Y. I. Li et al., 2016). We used RNA sequencing data collected in the same LCLs as used in this study. The data collection information can be found in Pickrell et al, 2010 and further processing can be found in Li et al. 2016 (Pickrell et al., 2010; Y. I. Li et al., 2016). We computed a nascent transcription phenotype for each gene as the normalized 4su expression level over the sum of the RNA expression and 4su expression level. We calculated the correlation between this value as well as nuclear 3’ Seq read counts for each gene divided by the sum of total 3’ Seq read counts and nuclear 3’ Seq read counts for each gene. We also calculated the correlation between the difference in the number of PAS identified both fractions and the nascent transcription phenotype using the summary lm function in R.

### apaQTL calling in both fractions

We used the leafcutter prepare phenotype table.py script with default settings to normalize the PAS usage ratios across individuals within each fraction. This method also outputs the top principal components (PCs) of the data to use as covariates. We plotted the proportion of variation explained by each PC in order to identify the number of PCs to include in the analysis (Supplementary Fig. 8). We included the top 4 PCs as well as the library preparation batch as the covariates. The top four PCs correlate most strongly with the cell count at collection (Supplementary Fig. 8). We used the same genotypes from Li et al. 2016(Y. I. Li et al., 2016), available at http://eqtl.uchicago.edu/jointLCL/genotypesYRI.gen.txt.gz (Y. I. Li et al., 2016). We removed individual NA19092 due to lack of genotype information in this file, bringing our sample size to 51 individuals for this part of the analysis. Only SNPs with a MAF > 5% in our sample were included. We used FastQTL to map apaQTLs in cis (25kb on either side) with 1000 permutations to select the top SNP-PAS association (Ongen, Buil, Brown, Dermitzakis, & Delaneau, 2016). We called apaQTLs in each fraction as variants passing 10% FDR (Benjamini-Hockberg) after permutations. In order to plot interpretable effect sizes for each association we computed nominal PAS:SNP associations for the pre-normalized PAS ratios.

### Association of apaQTLs with chromatin states

We downloaded the GM12878 chromatin HMM annotations for Hg19 from the UCSC table browser (Kent et al., 2002). We overlapped the eQTLs identified and published in Li et al. 2016(Y. I. Li et al., 2016) as well as the total and nuclear fraction apaQTLs with these categories. We calculated 95% confidence intervals for each measurement by sampling the number of QTLs in the set with replacement 1000 times (Fig. 3d and Supplementary Fig. 18).

### apaQTL overlap with eQTLs

We obtained the set of explained and unexplained eQTLs from Li et al. 2016 (Y. I. Li et al., 2016). In order to test whether genes with an unexplained eQTL are more likely to be explained by variation in APA, we separated the permuted apaQTL association (top snp per PAS) into three categories: unexplained eGene, explained eGene, non eGenes. We tested for significant enrichment of apaQTLs in each category using onesided Wilcoxon rank sum tests. In order to test if each explained and unexplained eQTLs described in Li et al. 2016(Y. I. Li et al., 2016) overlaps with an apaQTL, we extracted the nominal associations for each eQTL gene-SNP pair from the apaQTL data in both fractions. In order to account for multiple PAS associations for each pair, we selected the most significant p-value and used a Bonferroni correction to account for the number of PAS tested in the gene. We consider an eQTL as explained by an apaQTL if the corrected p-value is less than 0.05 but report the values for a range of cutoffs in (Supplementary Fig. 19).

### apaQTLs overlap with protein specific QTLs

The list of protein specific QTL genes can be found in the supplementary information from Battle et al. 2015(Battle et al., 2015). In order to show that genes with an eQTL and protein specific QTLs are likely to be associated with APA, we separated the permuted apaQTL association (top snp per PAS) into three categories: eGene, pGene, or neither pGene or eGene. We tested for significant enrichment with one sided Wilcoxon rank sum tests.

### Identification of molecular QTL associations

We sought to test if SNPs identified as apaQTLs are significantly associated with other molecular phenotypes previously tested in the same panel of LCLs. We tested for associations between the genotypes used in this study and each gene for each phenotype with fastqtl using the top 5 PCs calculated in Li et al. 2016 as covariates (Y. I. Li et al., 2016). We used normalized RNA expression, RiboSeq values, and protein levels, published in Li et al. 2016 (Y. I. Li et al., 2016).

### apaQTL overlap with GWAS Catalog

We downloaded the CRCh37hg19 GWAS catalog for UCSC table browser (Kent et al., 2002). We identified SNPs in LD with the nuclear apaQTLs using the LDproxy tool from LDlink with YRI as the population (Machiela & Chanock, 2015). We filtered all results to SNPs with an r2 greater than 0.9. We overlapped the full set with the GWAS catalog using pybedtools.

## Acknowledgements

We thank N. Gonzalez, J.P. Staley, M.C. Ward for comments on the manuscript.

## Funding

This work was supported by the US National Institutes of Health (R01GM130738 to Y.I.L) B.E.M. supported by T32 GM09197 to the University of Chicago and F31HL149259 to B.E.M. from National Heart, Lung, And Blood Institute of the National Institutes of Health. SP was in part supported by the National Center for Advancing Translational Sciences of the NIH (K12 HL119995). This work was completed in part with resources provided by the University of Chicago Research Computing Center.

## Author Contributions

Y.I.L. conceived of the project. B.E.M, S.W. and S.P. performed the experiments. B.E.M analyzed the data with help from Y.I.L, S.P., T.Z. and M.K. B.E.M. drafted the manuscript with input from Y.G., Y.I.L, and S.P. S.P., Y.G. and Y.I.L. supervised this project.

## Competing interests

The authors declare no competing interest.

## Data and material Availability

Fastq files and PAS annotations are available at GEO under accession GSE138197. All reproducible scripts can be found at https://brimittleman.github.io/apaQTL/

