## Supplemental Figures for "Alternative polyadenylation mediates genetic regulation of gene expression"

**Supplementary Figure 1: Relationship between 3' Seq and RNA decay, supplement to figure 1e**

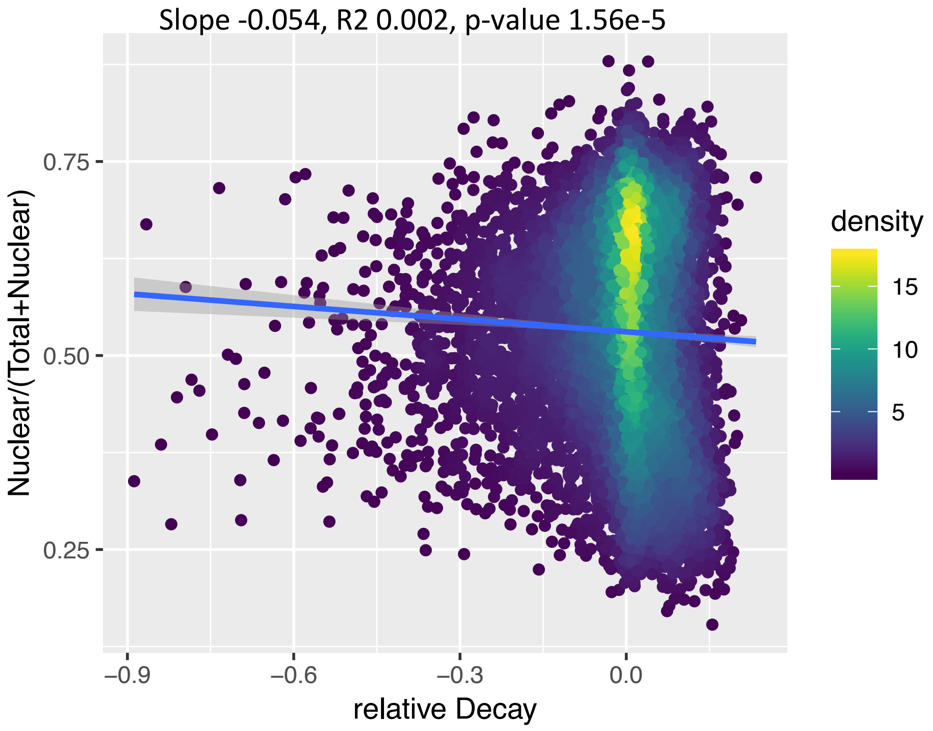


Correlation between the ratio of nuclear to total reads and relative RNA decay rate for each gene. RNA Decay data obtained from Pai et al. [1]

**Supplementary Figure 2: Distribution of signal site in sequence upstream of PAS. Supplement to Fig. 1c.**

a b

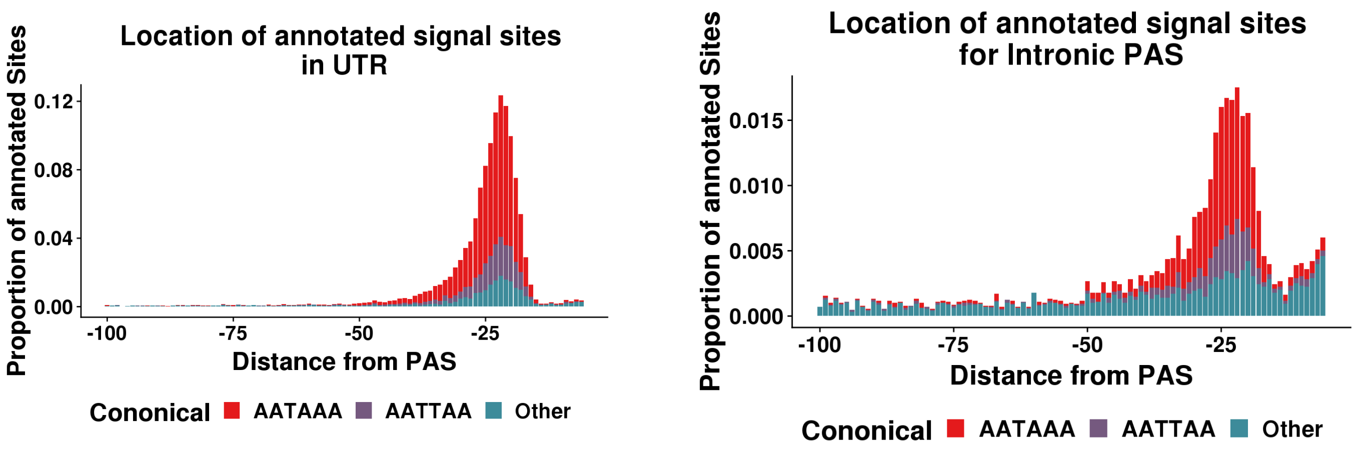


a. Signal site distribution for PAS in 3' UTR. Signal site sequences in "Other" shown on figure 1c.

b. Signal site distribution for Intronic PAS. Signal site sequences in "Other" shown on figure 1c.

**Supplementary Figure 3: Proportion of PAS in each genic location for total fraction. Additional figures corresponding to figure 1c.**

a

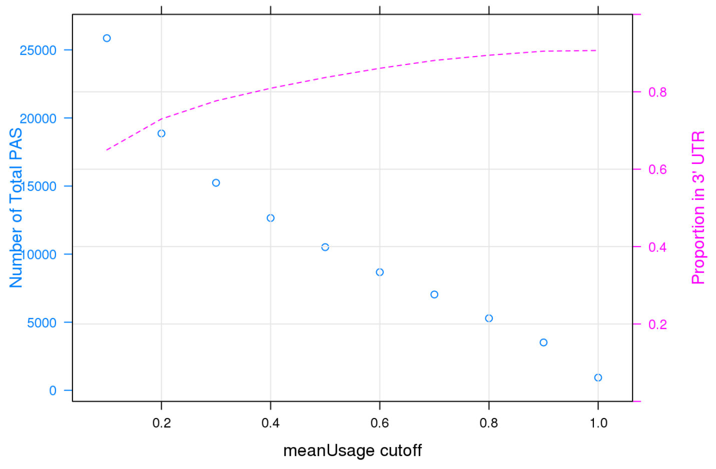


b

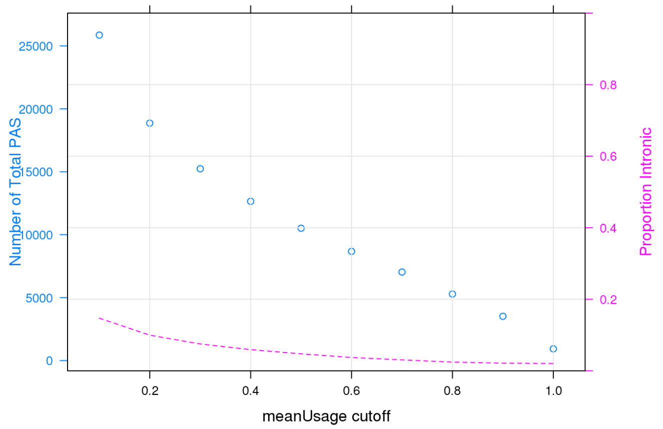


a. Number of PAS and proportion of PAS in the 3' UTR according to mean usage in the total fraction.
b. Number of PAS and proportion of PAS in introns according to mean usage in the total fraction.

**Supplementary Figure 4: Location of PAS differentially used**

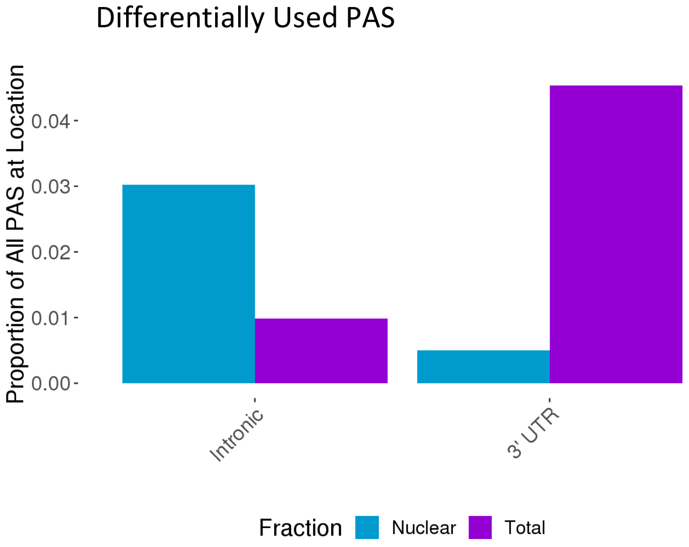


We used the *leafcutter* to identify genes with significant differential usage of PAS between the total and nuclear fraction. The majority of PAS preferentially used in the nuclear fraction are intronic and the majority of PAS preferentially used in the total fraction lie in the 3' UTR.

### Supplementary Figure 5. Comparison to previous PAS annotations

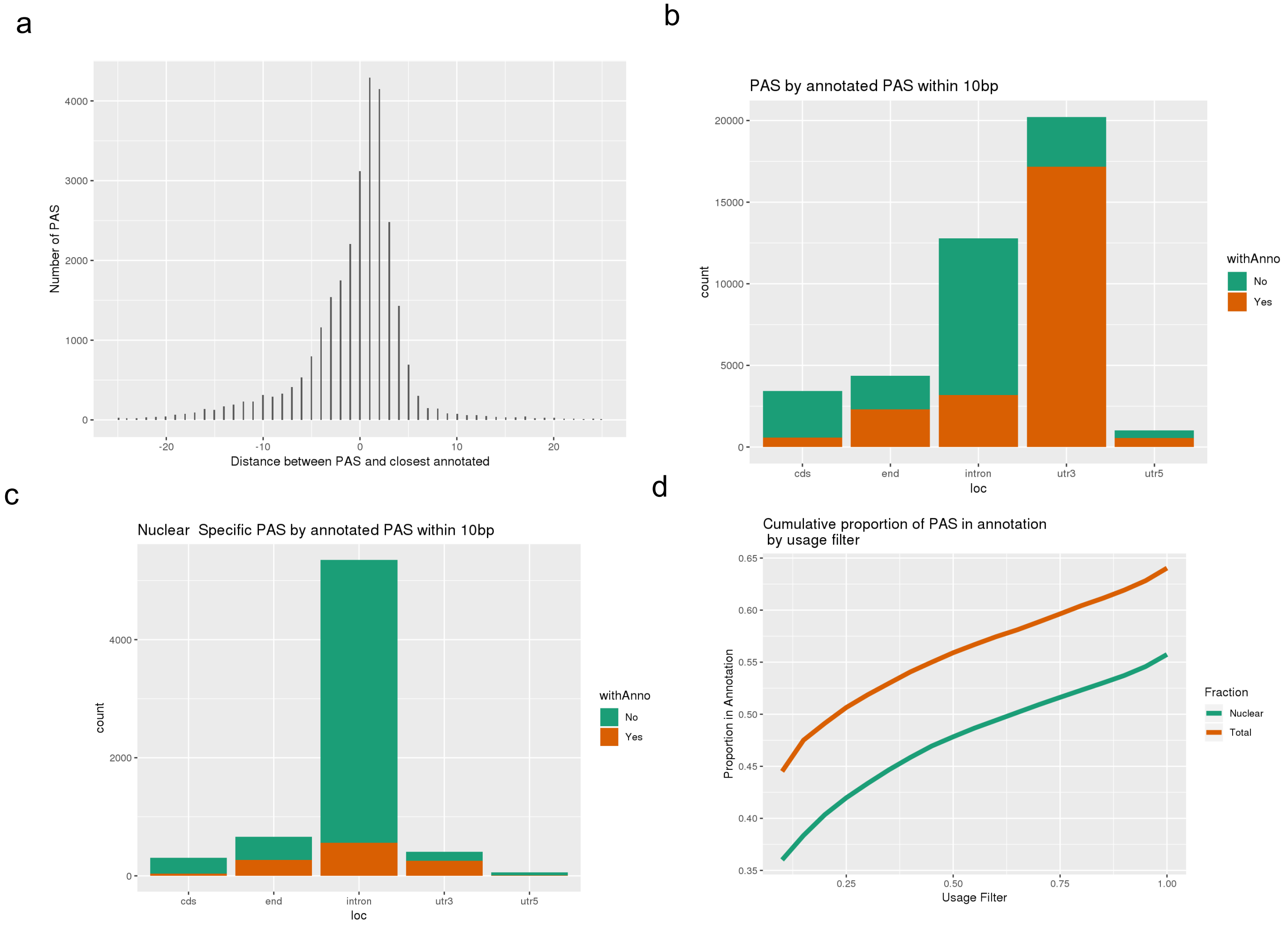


a. Distance between PAS and closest annotated site in the annotation database (PolyA_DB release 3.2)[2].
b. PAS at each genic location according to presence of annotated PAS within 10bp.
c. Nuclear specific PAS by location. Colored by presence in annotation within 10pb.
d. Proportion of PAS in annotation by usage in each fraction.

**Supplementary Figure 6: Q-Q plots for apaQTLs**

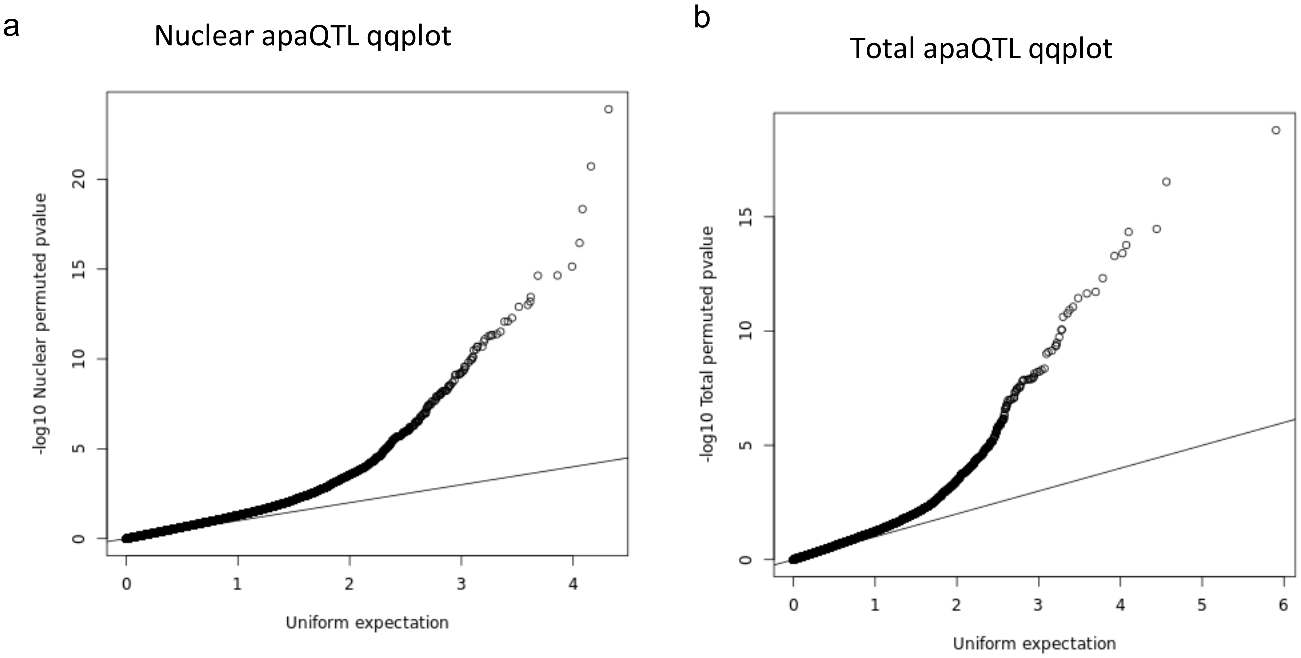


a. Q-Q plot for Nuclear apaQTLs, plotting top SNP PAS associations.
b. Q-Q plot for Total apaQTLs, plotting top SNP PAS associations.

**Supplementary Figure 7: Proportion of PAS tested with an apaQTL**

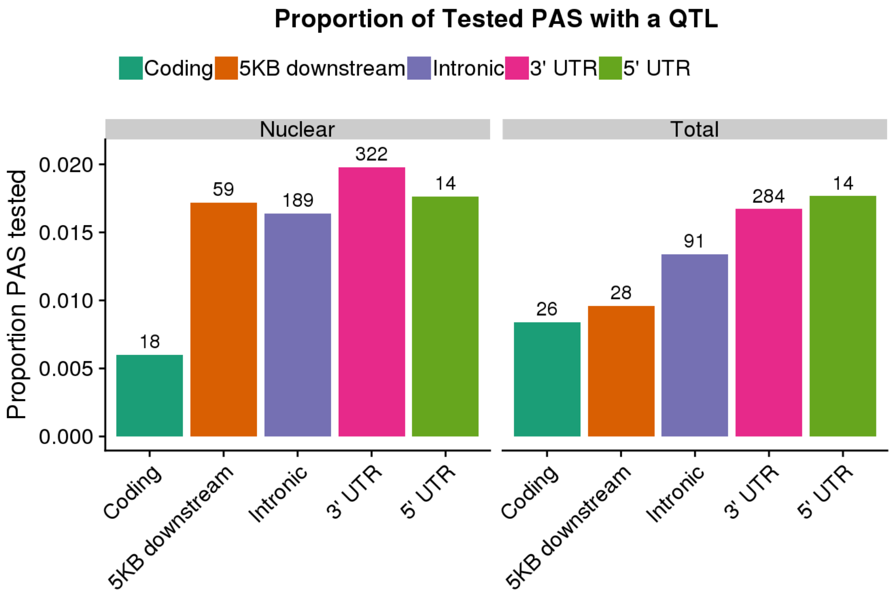


Proportion of PAS in each location of those tested with a significant apaQTL. Numbers represent number of identified apaQTL in given location.

### Supplementary Figure 8: Evaluation of number of PCs for apaQTL analysis

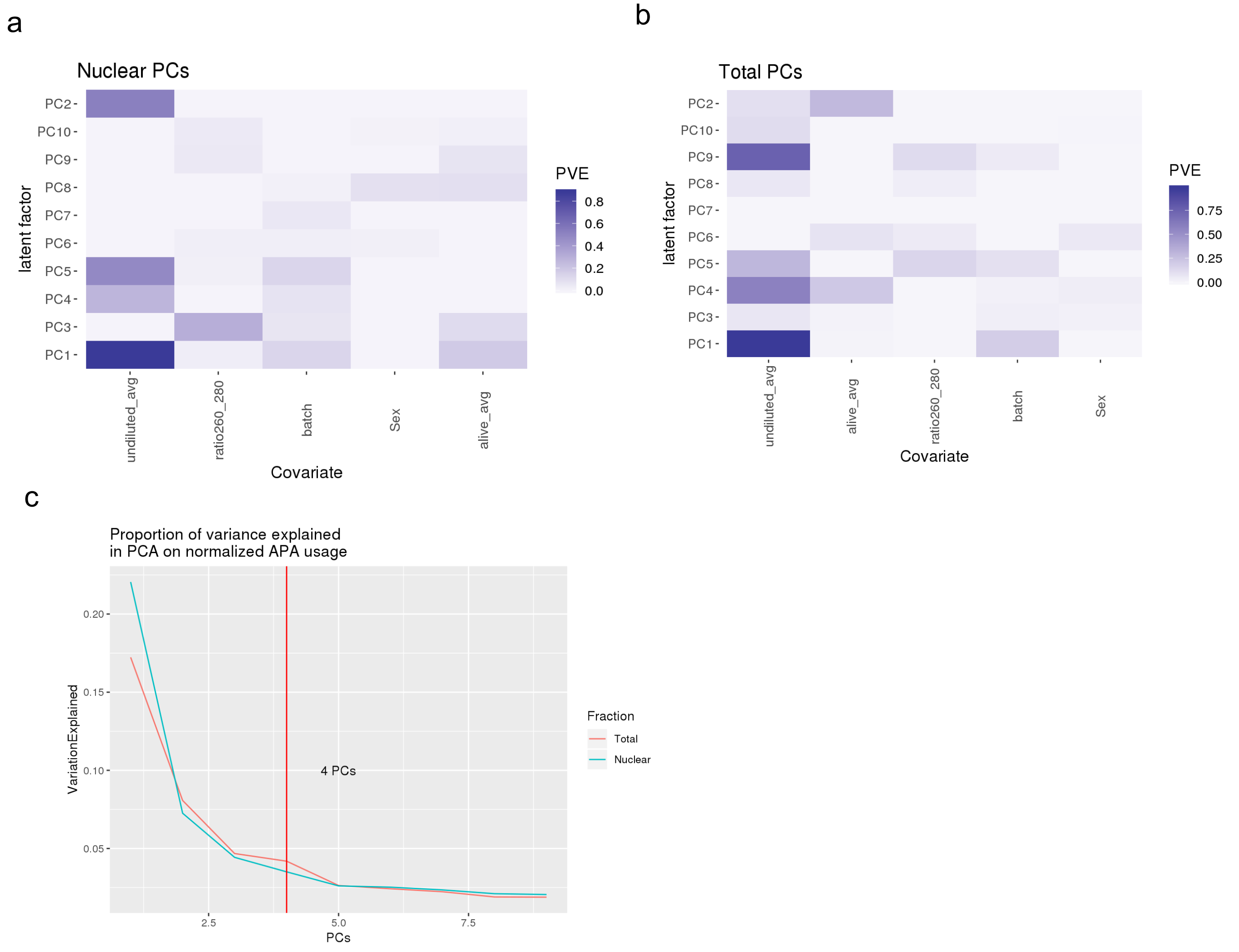


a. Proportion of variance explained in each PC by experimental variables in nuclear APA phenotype
b. Proportion of variance explained in each PC by experimental variables in total APA phenotype
c. Proportion of variance explained by each PC in APA phenotypes. Vertical line used to denote number of PCs accounted for in apaQTL analysis.

### Supplementary Figure 9: Signal site disruption

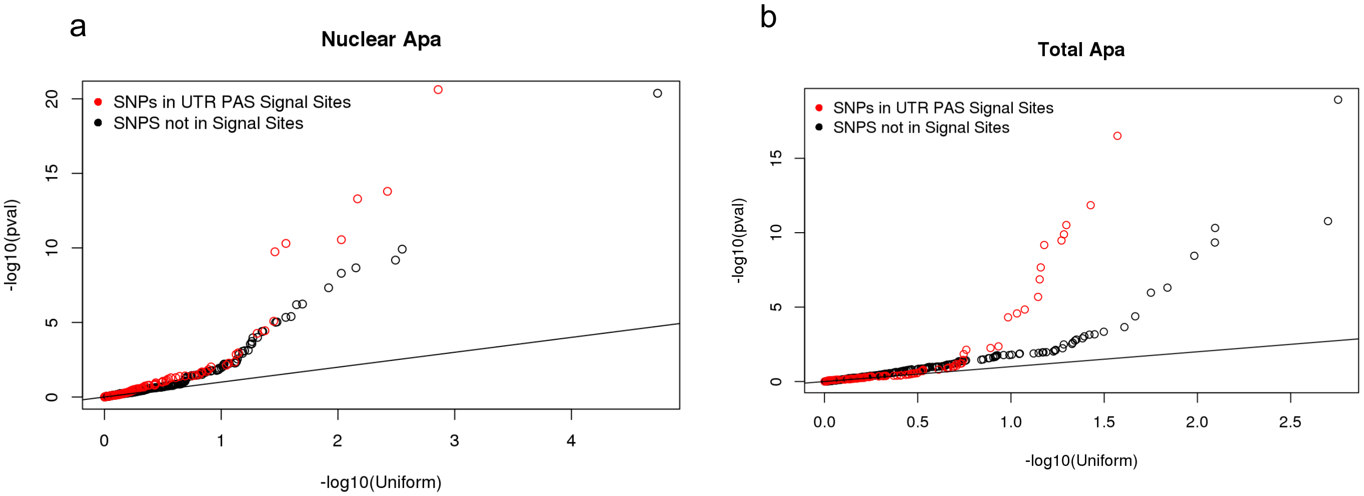


a. Nuclear apaQTL p-values for SNP in signal sites upstream of 3' UTR PAS compared to SNPs equal distance upstream of a set of 3'UTR PAS without identified signal sites upstream.
b. Total apaQTL p-values for SNP in signal sites upstream of 3' UTR PAS compared to SNPs equal distance upstream of a set of 3'UTR PAS without identified signal sites upstream.

**Supplementary Figure 10: apaQTL sharing between fractions.**

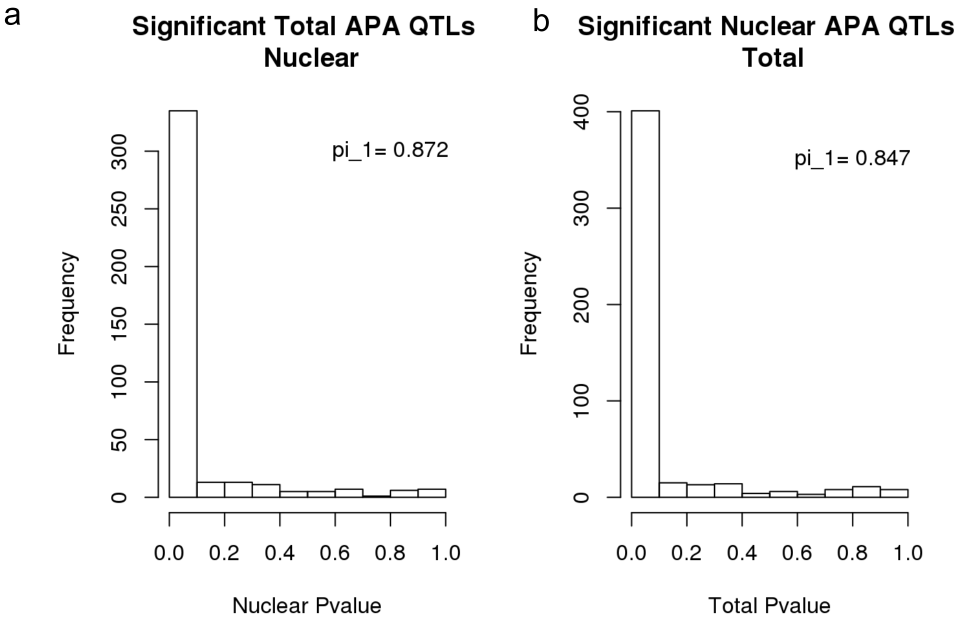


a. P-value distribution for Total apaQTLs in nuclear fraction. Values were calculated based on PAS tested in both fractions (403 of 443). Results are robust to using all PAS (pi_1=0.842)
b. P-value distribution for Nuclear apaQTLs in the total fraction. Values calculated based on PAS tested in both fractions. (483 of 602) Results are robust to using all PAS (pi_1=0.825)

### Supplementary Figure 11: Correlation of effect size

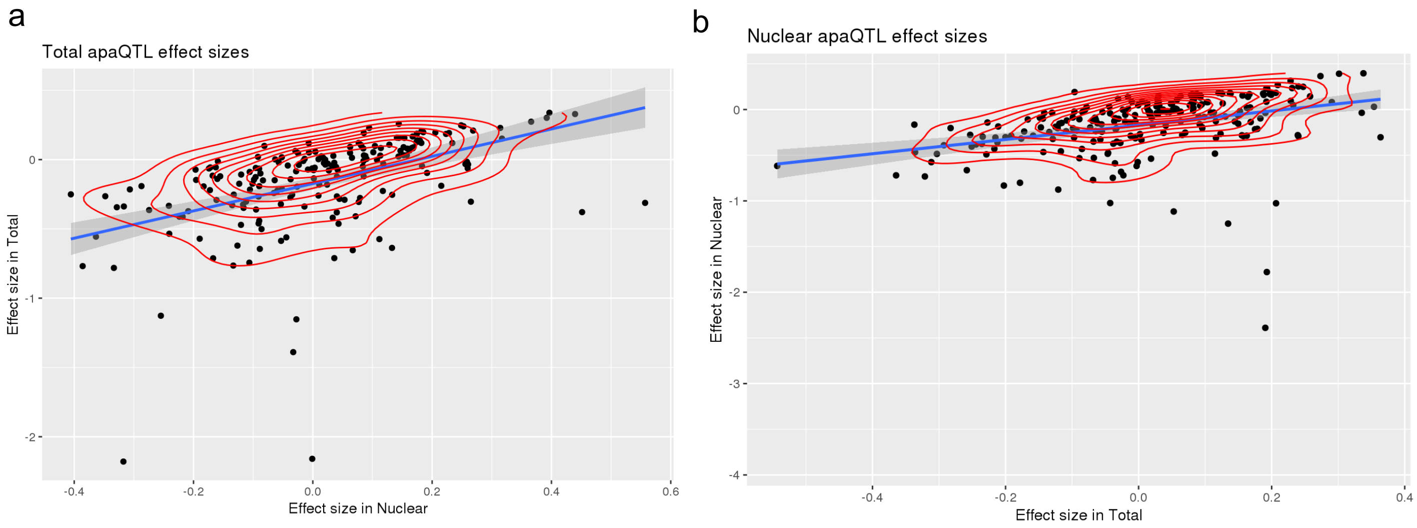


a. Normalized effect sizes in total and nuclear fraction for total apaQTLs tested in both fractions.
b. Normalized effect sizes in total and nuclear fraction for nuclear apaQTLs tested in both fractions.

**Supplementary Figure 12: Overlap between apaQTLs in total fraction and other molecular QTLs, supplement to Figures 3a and 3c.**

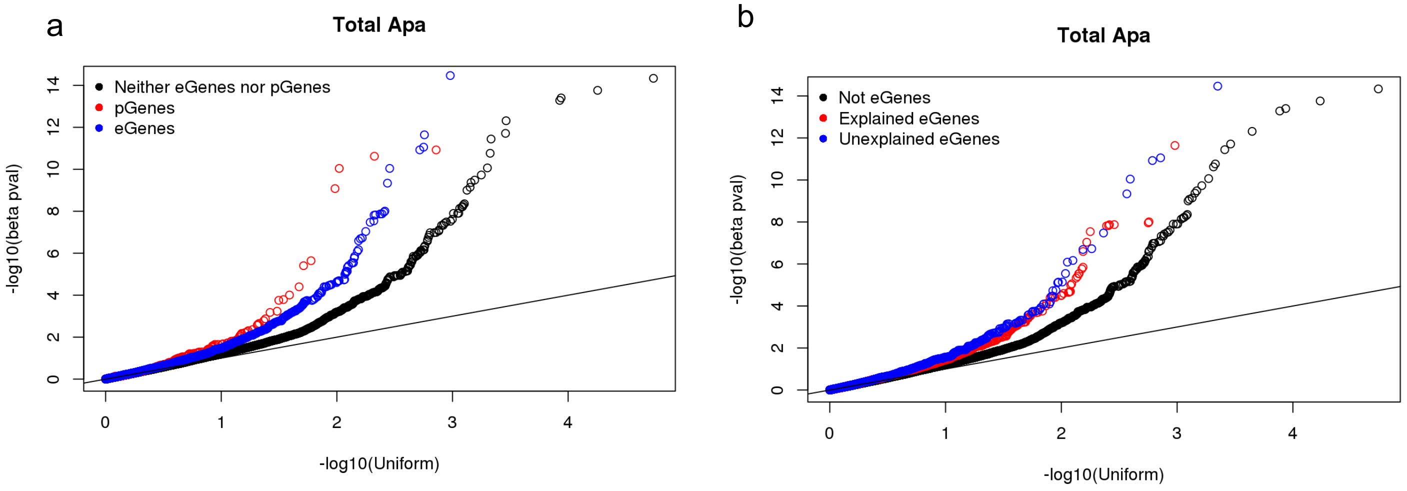


a. Total apaQTL permuted p-values separated by presence of pQTL, eQTL or neither type of QTL in the gene. Enrichment for low apaQTL p-values in genes with and eQTL or pQTL.
b. Total apaQTL permuted p-values separated by presence of explained or unexplained eQTLs. Enrichment for low apaQTL p-values in genes with explained eQTL and unexplained eQTL.

**Supplementary Figure 13: Figure 3b without outlier SNP**

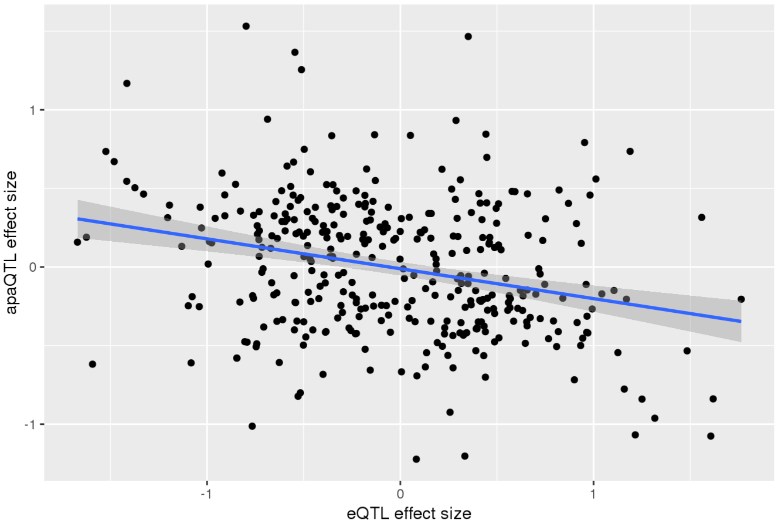


Relationship between intronic nuclear apaQTL effect size and eQTL effect size is not driven by outlier SNP in Figure 3b. Filtered for SNPs with eQTL effect size > -2.

**Supplementary Figure 14: Overlap between apaQTLs in total fraction and other molecular QTLs, supplement to Figures 3a and 3c.**

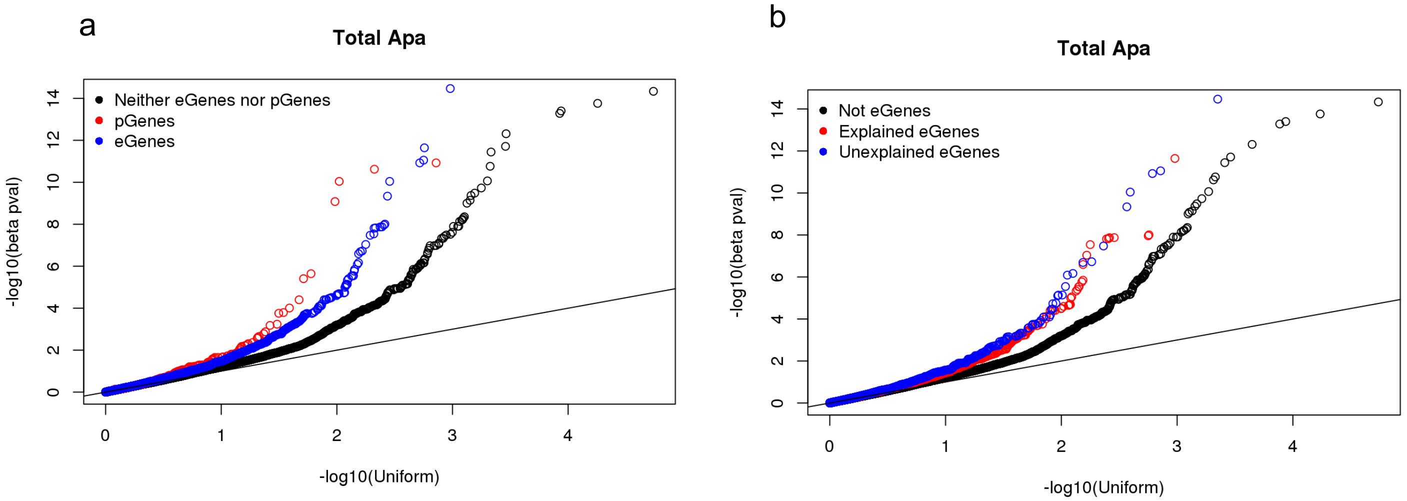


a. Total apaQTL permuted p-values separated by presence of pQTL, eQTL or neither type of QTL in the gene. Enrichment for low apaQTL p-values in genes with and eQTL or pQTL.
b. Total apaQTL permuted p-values separated by presence of explained or unexplained eQTLs. Enrichment for low apaQTL p-values in genes with explained eQTL and unexplained eQTL.

**Supplementary Figure 15: Locus Zoom Plots *EIF2A*, supplement to figure 4b**

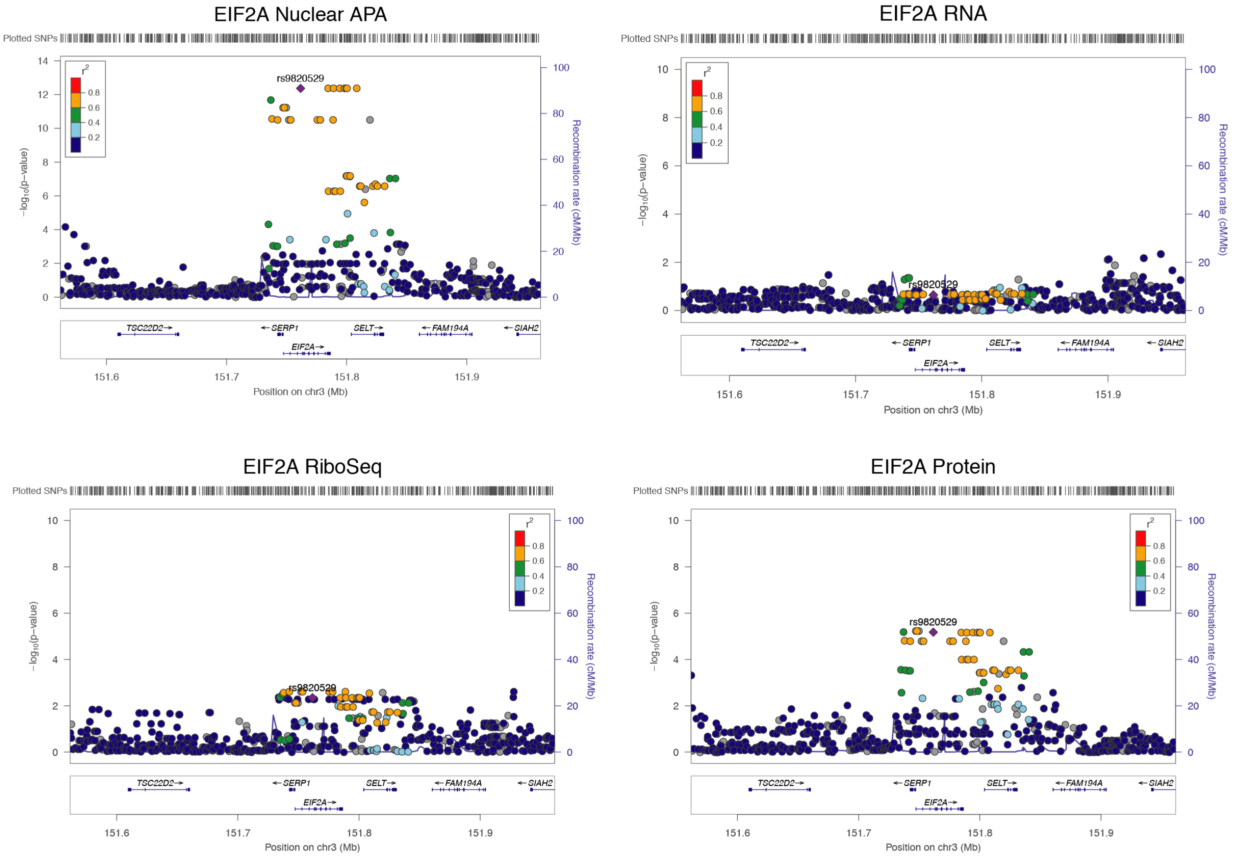


Locus zoom plots for *EIF2A* apaQTL in figure 4b. Associations in RNA, riboseq, and protein determined using normalized data from Li et al. 2016[2]. LD patterns according to HapMap YRI lines. Generated with Locuszoom.org.

**Supplementary Figure 16: Western Blots to demonstrate cell fractionation**

a

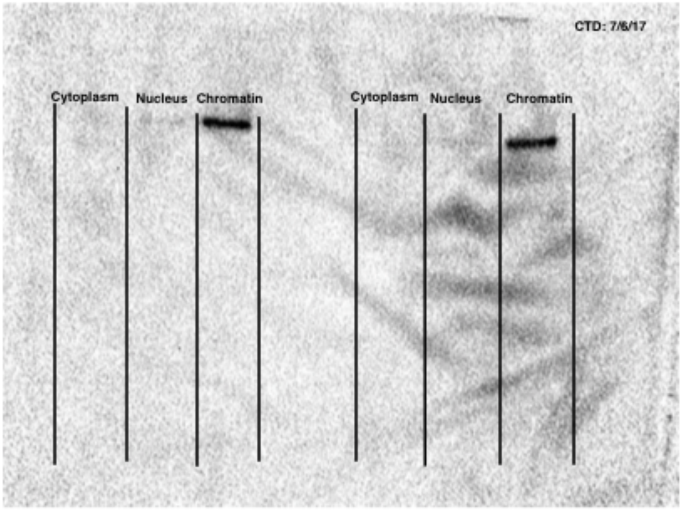


b

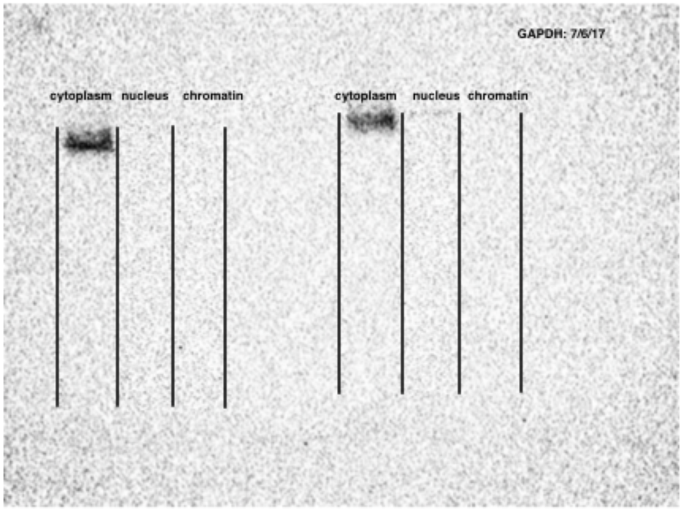


a. Western blot against Carboxyl terminal domain of RNA Polymerase II, photo captured at 10 second exposure. Blot is not used for quantification, but to validate cell fractionation.
b. Western blot against GAPDH to mark glycolysis in cytoplasm, photo captured at 25 second exposure time. Blot is not used for quantification, but to validate cell fractionation. Figure panels are modeled off Mayer and Churchman 2016, Figure 2[3].

### Supplementary Figure 17: 3' RNA-seq read mapping

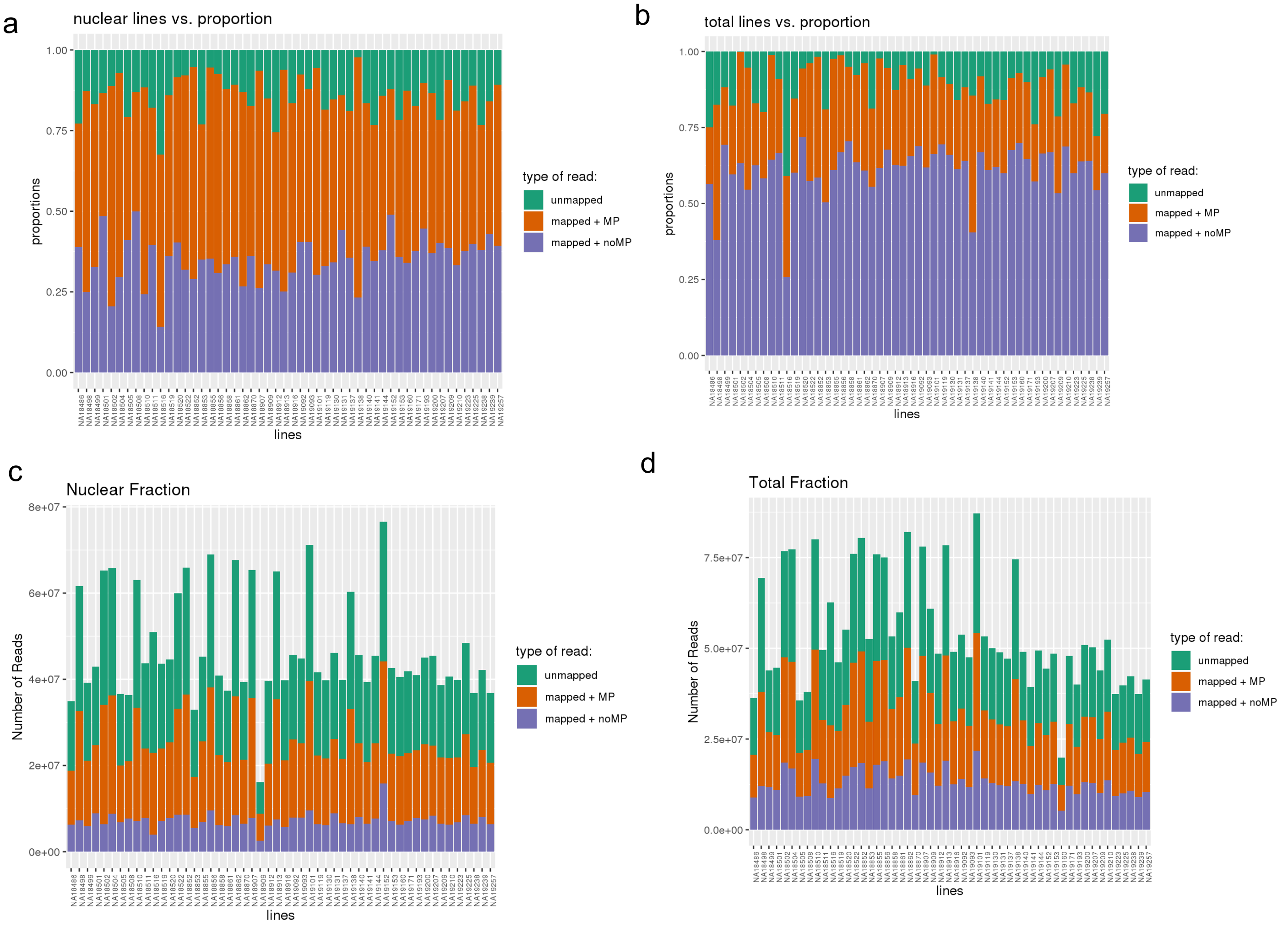


a. Proportion of reads per line in nuclear fraction, final reads used for analysis are mapped + noMP. These reads have mapped and passed the filtering for mispriming (MP) events.
b. Number of reads per line in nuclear fraction, final reads used for analysis are mapped + noMP. These reads have mapped and passed the filtering for mispriming events.
c. Proportion of reads per line in total fraction, final reads used for analysis are mapped + noMP. These reads have mapped and passed the filtering for mispriming events.
d. Number of reads per line in total fraction, final reads used for analysis are mapped + noMP. These

**Supplementary Figure 18: Location of apaQTLs and eQTLs by chromHMM category. Supplement to Figure 3d.**

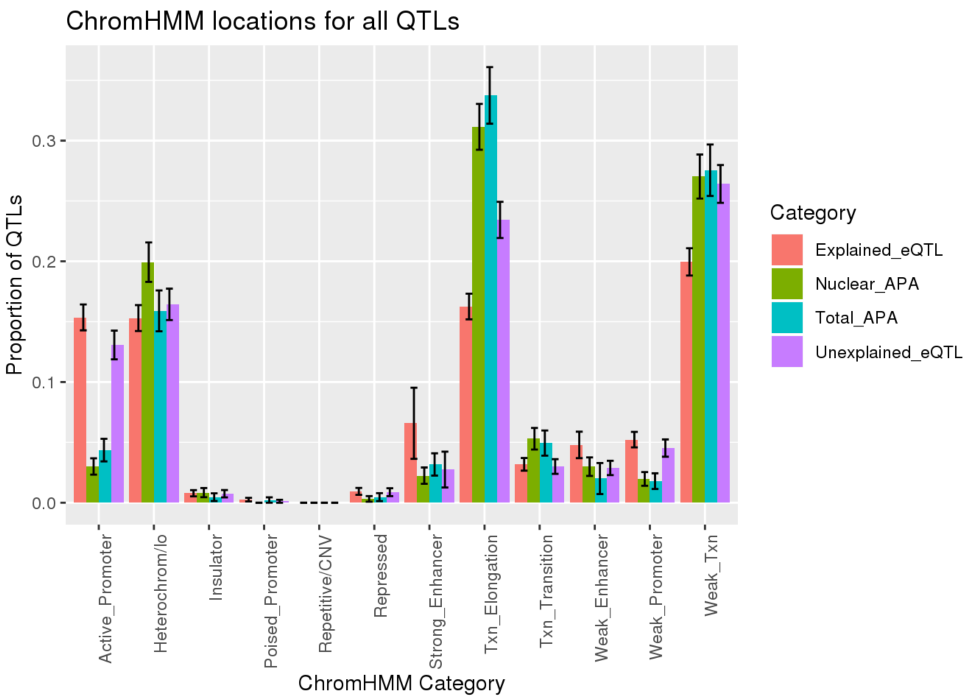


Proportion of QTLs falling in each of the 12 chromHMM categories. Error bars represent the 95% confidence interval for each measurement by bootstrapping 1000 times.

**Supplementary Figure 19: Proportion of eQTLs explained by apaQTLs**

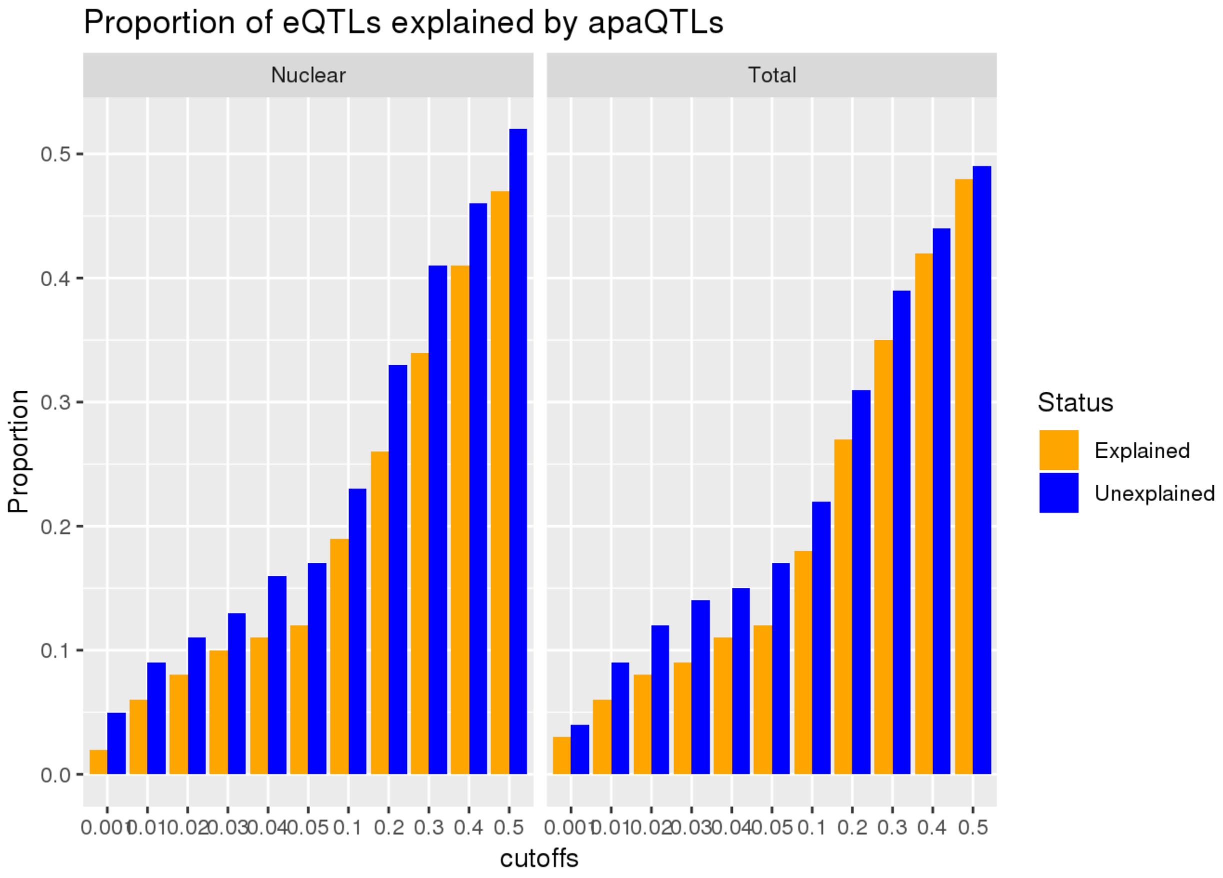


,

Proportion of eQTLs explained by apaQTLs separated by fraction. In both fractions, robust to different apaQTL p-value cutoffs. apaQTLs explain a higher proportion of previously unexplained eQTLs. Status comes from Li et al. 2016[2].

**Supplementary Table 1: ExpressionIndependentapaQTLs.txt**

apaQTL nominally significant in protein and not expression. Includes pvalue and slope for nuclear apa, expression, protein and riboseq data.

**Supplementary Table 2: MetaDataSequencing.txt**

Library information for each line. Sample, Collection, Read information
